# Optimal resistance management for mixtures of high-risk fungicides: robustness to the initial frequency of resistance and pathogen sexual reproduction

**DOI:** 10.1101/2022.02.14.480407

**Authors:** Nick P Taylor, Nik J Cunniffe

## Abstract

There is a strong consensus that selection for fungicide resistant pathogen strains can be most effectively limited by using applications of mixtures of fungicides designed to balance disease control against selection. However, how to do this in practice is not entirely characterised. Previous work indicates optimal mixtures of pairs of fungicides which are both at a high risk of resistance can be constructed using pairs of doses which select equally for both single resistant strains in the first year of application. What has not been addressed thus far is the important real-world case in which the initial levels of resistance to each fungicide differ, for example because the chemicals have been available for different lengths of time. We show how recommendations based on equal selection in the first year can be sub-optimal in this case. We introduce a simple alternative approach, based on equalising the frequencies of single resistant strains in the year that achieving acceptable levels of control is predicted to become impossible. We show that this strategy is robust to changes in parameters controlling pathogen epidemiology and fungicide efficacy. We develop our recommendation using a pre-existing, parameterised model of *Zymoseptoria tritici* (the pathogen causing Septoria leaf blotch on wheat), which exemplifies the range of plant pathogens which predominantly spread clonally, but for which sexual reproduction forms an important component of the life cycle. We show that pathogen sexual reproduction can influence the rate at which fungicide resistance develops, but does not qualitatively affect our optimal resistance management recommendation.

## Introduction

World food security faces multiple threats, including the growing global population (***Godfray et al., 2010***), climate change (***Tai et al., 2014***) and plant disease (***Strange and Scott, 2005***). However, it is estimated that food production will need to increase by 60% by 2050 (***Ristaino et al., 2021***). Despite annual spending of roughly 16 billion US dollars on fungicides globally, estimated crop losses due to disease stand at 20% (***Jorgensen et al., 2017***). Fungicide resistance challenges our ability to maintain control of fungal pathogens, but effective resistance management strategies prolong control of these yield-limiting crop diseases and have been studied for decades (***Staub, 1991***; ***van den Bosch et al., 2014a***; ***Corkley et al., 2021***). We explore the optimal management of fungicide resistance in crop pathogens using mixtures containing pairs of fungicides which are ‘high-risk’ for resistance. In particular, we explore the effects of sexual pathogen reproduction and variation in initial levels of resistance.

We use Septoria leaf blotch (*Zymoseptoria tritici*), the most prevalent disease of wheat worldwide (***Suffert et al., 2011***), as our case study. An estimated 70% (≈€1bn) of the European cereal fungicide market is primarily targeted towards the management of *Z. tritici* of winter wheat (***Torriani et al., 2015***). Most fungicide modelling studies focus on Septoria, so there are existing parameterised models available (e.g. ***Hobbelen et al***. (***2013***); ***Elderfield et al***. (***2018***)). Further, it is a heterothallic fungus (***Suffert et al., 2016***) capable of both sexual and asexual reproduction (***Suffert et al., 2011***; ***Eriksen et al., 2001***; ***Singh et al., 2021***). It therefore exemplifies the large number of plant pathogens for which sexual reproduction is potentially important in the epidemiology (***Agrios, 2004***) and evolution (***McDonald et al., 1996***). For Septoria, reported proportions of sexual reproduction differ widely between experiments (***Chen and McDonald, 1996***; ***Zhan et al., 1998***), but it is known that ascospores produced via sexual reproduction initiate Septoria epidemics within a field (***Shaw and Royle, 1989***). Although ascospores are quantitatively the most significant form of primary inoculum (***Suffert et al., 2011***), for simplicity most fungicide resistance modelling studies do not consider pathogen sexual reproduction, although there are some exceptions (***Shaw, 1989***). Here we seek to understand the effect of the inclusion of pathogen sexual reproduction on the resulting resistance management recommendation. We neglect the effect of ascospores within the growing season, due to previous studies suggesting they have a small effect on the severity of an epidemic (***Eriksen et al., 2001***), in part caused by the longer latent period of the sexual pseudothecia compared to the clonal pycnidia. Optimal management principles for Septoria may transfer to other fungal and oomycete crop pathogens which reproduce sexually, e.g. *Phytophthora infestans*, cause of the potato disease late blight (***Fones et al., 2020***).

A common resistance management strategy is to use fungicide mixtures with more than one mode of action present in the mixture. These fungicides are often categorised as ‘low-risk’ or ‘high-risk’ for resistance, depending on whether resistant pathogen strains exist in the population and the likelihood of developing resistance, amongst other factors (***Brent and Hollomon, 2007***). In practice fungicide mixtures often contain two fungicides that are high-risk for development of resistance. These mixtures are of increasing relevance since there are few low-risk fungicides available and the high risk options are typically of higher efficacy (***van den Bosch et al., 2014b***). Further, lowrisk (multi-site) fungicides are increasingly rare; for example chlorothalonil has been banned for use by the EU due to environmental concerns since 2019 (***Murray, 2019***). Previous modelling studies have found fungicide mixtures to be more effective as a resistance management strategy than alternating use of fungicides (***Elderfield et al., 2018***) or spatially concurrent applications (***Hobbelen et al., 2013***), where different fields receive treatments from different modes of action. That mixtures outperformed alternations or concurrent use was robust to fitness costs, partial resistance, changes in fungicide parameters and the initial frequency of the double resistant strain. For this reason we concentrate exclusively here on the optimal strategy for mixtures of two high-risk fungicides and seek to test how to optimally construct high-risk fungicide mixtures if current levels of resistance to the two mixing partners differ, or if between-season pathogen sexual reproduction is considered.

It was reported by ***van den Bosch et al***. (***2014b***) that, across 17 publications, mixtures of high-risk fungicides resulted in a reduction in selection for resistance in 20 out of 24 pathogen-crop-fungicide combinations. There is ongoing debate about how high-risk mixtures (i.e. mixtures of high risk fungicides) should be constructed. Although the so-called ‘governing principles’ (***van den Bosch et al., 2014a***) suggest that increased fungicide dose increases selection for a given mode of action, increased dose of a mixing partner can reduce selection for the other mode of action in the mixture (***van den Bosch et al., 2014b***). Modelling work shows that the optimal way to mix a low-risk and a high-risk fungicide is to use the maximum dose of the low-risk chemical and the minimal viable dose of the high-risk chemical (***Hobbelen et al., 2011a***; ***Elderfield et al., 2018***). However, maximising the dose of either fungicide when the mixture contains two high-risk chemicals could lead to excessive selection pressure on that fungicide, so different recommendations are required.

***Hobbelen et al***. (***2013***) use modelling to address the case where the fungicide mixture contains two high-risk chemicals. They consider four pathogen strains: one that is resistant to both chemicals; one that is sensitive to both; and two more that are sensitive to one fungicide but resistant to the other. Their results suggest that the choice of doses used is crucial to the resulting durability of the strategy. ***Hobbelen et al***. (***2013***) suggest the optimal fungicide mixture has a dose pairing that is as weak as possible whilst achieving sufficient yield, and selects equally for both single resistant strains. These authors addressed the case where both single resistant strains are initially at the same frequency, and explored what happens for different amounts of the double resistant strain. However, that study did not address the common real-world scenario where the initial levels of resistance to the two chemicals differ. The optimal strategy in this case is not yet described and – as well as the effect of sexual reproduction – is the focus of this paper. In this work we introduce and test a new prescription based on equalising the resistance frequencies by the time of breakdown, rather than equalising selection in the first year.

Initial levels of resistance commonly differ for different fungicides due to differing natural incidences of resistant strains, or because one fungicide was introduced to market much earlier than its mixing partner. For instance, resistance to benzimidazoles developed rapidly in the mid-1980s (***Blake et al., 2018***), but resistance to Quinone outside Inhibitors (QoI) fungicides was not detected in the UK until 2001 (***Cheval et al., 2017***). Benzimidazoles were introduced in the 1960s, but QoIs were not introduced until the 1990s (***Leadbeater, 2014***), suggesting levels of resistance and rate of increase of resistance differed greatly between these fungicide classes. Fungicide sensitivity has been reported to differ for high-risk methyl benzimidazole carbamate (MBC) fungicides depending on year and region, with sensitive proportions of *Oculimacula acuformis* (eyespot disease of cereals) comprising 92% in Germany in 1985, but only 4% and 16% in France and the UK respectively in the same year (***Parnell et al., 2008***). Differing resistance frequencies in *Botrytis cinerea* (gray mould of raspberries) to seven different fungicides from a variety of fungicide classes were reported in Northern Germany (***Rupp et al., 2017***). Further, initial resistance frequencies may be influenced by mutation-selection balance (***van den Bosch and Gilligan, 2008***) which depends on the fitness costs of resistance, which will depend on the mutation, and hence the fungicide.

Fungicides can be described by ‘dose-response curves’ which are measures of their efficacy. These curves differ depending on the mode of action and effectiveness of each chemical. We seek to show how different dose-response parameters influence the optimal strategy even when levels of resistance to the two fungicides vary and/or pathogen sexual reproduction is present. Understanding the effect of different dose response curves on the optimal strategy is crucial to decision making when constructing mixtures containing pairs of existing fungicides and/or new chemicals that come on to the market. These decisions are made by agronomists and growers but typically informed by recommendations from the Fungicide Resistance Action Committee (FRAC).

In this paper we address the following questions:

1. What is the effect of varying initial levels of resistance on optimal resistance management strategies for mixtures of pairs of high risk fungicides?
2. When do existing strategy recommendations fail, and how can we improve upon them?
3. How robust is our new recommendation to alterations in parameters controlling pathogen epidemiology and fungicide efficacy?
4. What is the effect of the balance of between-season sexual and asexual reproduction on the model and the strategy recommendation?

## Methods

The model is an adapted version of one presented by ***Hobbelen et al***. (***2013***), which addresses two high-risk fungicides used together to control Septoria. The model is compartment-based and measures different categories of leaf tissue (Figure 2). After infection, healthy (susceptible) tissue (*S*) transitions to exposed tissue (*E*) (infected but not infectious) and then to infectious tissue (*I*), before removal (*R*) – see Figure 2. The initial infection is given by a primary inoculum (*P*). The model also includes growth and senescence of living tissue. The model also tracks the active concentration of both fungicides in the mixture over time. We split the modelled year into two distinct time periods – within and between growing seasons. A full list of model parameters and values (values taken from ***Hobbelen et al***. (***2013***)) is provided in Table 1.

**Table 1.** Parameters and state variables used in the HRHR model. Sources: ***Hobbelen et al***. (***2013***); ***Elderfield et al***. (***2018***). Although the default value for the proportion of between-season sexual reproduction (*q*_*B*_) is 0, we test different values to explore the effect of this parameter on the results of the model. The value of *1/*_0_ is changed by the non-dimensionalisation process, and the function used by ***Hobbelen et al***. (***2013***) was *r*(*k* − *A*) for *k* = 4.2, not *r*(1 − *A*). All other parameter values are unchanged by this process. This explains why the tissue state variables (and some of the other parameters) have no units in our version of the underlying model.

| Symbol | Meaning | Type | Default value/range/equation | Units |
| --- | --- | --- | --- | --- |
| $S(t)$ | Susceptible tissue | Variable | [0, 1] | - |
| $E_{mn}(t)$ | Latently infected (exposed) tissue, strain $mn$ | Variable | [0, 1] | - |
| $I_{mn}(t)$ | Infectious tissue, strain $mn$ | Variable | [0, 1] | - |
| $R(t)$ | Removed tissue | Variable | [0, 1] | - |
| $A(t)$ | Total tissue ($= S + R + \sum_{m,n} [E_{mn} + I_{mn}]$) | Variable | [0, 1] | - |
| $P_{mn}(t)$ | Primary inoculum, strain $mn$ | Variable | $[0, \psi_0]$ | - |
| $C_i(t)$ | Fungicide $i$ concentration | Variable | $\geq 0$ | Label dose (fraction of) |
| $t$ | Time | Variable | $\geq 0$ | degree-days |
| $g(A)$ | Production of host leaf tissue | Function | Equation 7 | - |
| $\Gamma(t)$ | Senescence | Function | Equation 8 | - |
| $\delta_{F,m}(C_F)$ | Fungicide $F$ growth rate factor, strain $m$ | Function | Equation 9 | - |
| $r$ | Host growth rate | Parameter | $1.26 \times 10^{-2}$ | degree-days<sup>-1</sup> |
| $\beta$ | Infection rate | Parameter | $1.56 \times 10^{-2}$ | degree-days<sup>-1</sup> |
| $\gamma^{-1}$ | Latent period | Parameter | 266 | degree-days |
| $\mu^{-1}$ | Infectious period | Parameter | 456 | degree-days |
| $\psi_0$ | Initial inoculum amount | Parameter | $(1.09/4.2) \times 10^{-2}$ | - |
| $\nu$ | Inoculum decay rate | Parameter | $8.5 \times 10^{-3}$ | degree-days<sup>-1</sup> |
| $q_B$ | Between-season sexual reproduction proportion | Parameter | 0 | - |
| $\Lambda_F$ | Fungicide $F$ decay rate | Parameter | $1.11 \times 10^{-2}$ | degree-days<sup>-1</sup> |
| $\theta_F$ | Fungicide $F$ curvature | Parameter | 9.6 | Label dose<sup>-1</sup> |
| $\omega_F$ | Fungicide $F$ asymptote | Parameter | 1 | - |
| $T_{\text{emerge}}$ | Start of modelled season | Parameter | 1212 | degree-days |
| $T_{GS32}$ | First treatment time | Parameter | 1456 | degree-days |
| $T_{GS39}$ | Second treatment time | Parameter | 1700 | degree-days |
| $T_{GS61}$ | Senescence begins | Parameter | 2066 | degree-days |
| $T_{GS87}$ | End of season | Parameter | 2900 | degree-days |

We generalise the model presented by ***Hobbelen et al***. (***2013***) by introducing between-season sexual reproduction, because Septoria’s sexual ascospores are reported to contribute to a large proportion of the primary inoculum that initialises each epidemic (***Eriksen et al., 2001***; ***Suffert et al., 2011***). We denote the proportion of between-season sexual reproduction *q*_*B*_, and initially set *q*_*B*_ = 0 in line with ***Hobbelen et al***. (***2013***) before exploring the effect of between-season sexual reproduction by scanning over a range of values. We neglect within-season sexual reproduction for simplicity and because previous research suggests its effect on epidemic severity is small (***Eriksen et al., 2001***). Another change to the model presented by ***Hobbelen et al***. (***2013***) is that we consider one field instead of two, since we omit the less effective ‘concurrent field’ strategy.

### Pathogen strains

When studying fungicide resistance evolution, modellers often consider an emergence and a selection phase separately (***van den Bosch and Gilligan, 2008***; ***Milgroom, 1990***; ***van den Bosch et al., 2011***). The former concerns the initial stochastic phase where new resistant strains appear through random mutation and invasion. We do not account for the emergence phase, and focus entirely on the selection phase, which is where the resistant strain is already established in the population, and a selection pressure is applied when fungicide treatments are used.

We label the two fungicides *A* and *B*. It is assumed that there are four pathogen strains; the double sensitive strain, two single resistant strains and the double resistant strain. These are denoted by *ss, sr, rs* and *rr* respectively, where *r* indicates resistant and *s* indicates sensitive to fungicide *A* or *B* (for example the labelling *rs* would correspond to a pathogen strain that is resistant to fungicide *A* but sensitive to fungicide *B*).

### Within-season

#### Within-season model equations

The within-season model dynamics are as follows:

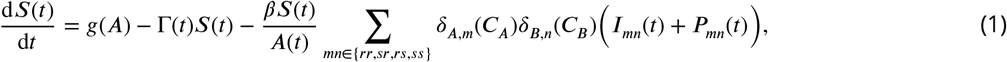

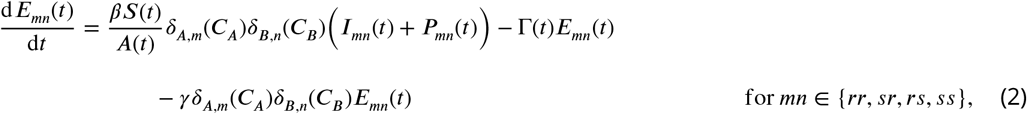

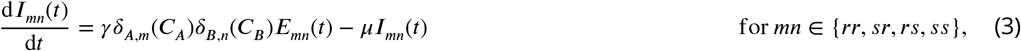

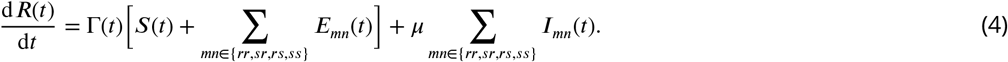

See Table 1 for parameter, variable and function definitions. Note that the notation *mn* ∈ {*rr, sr, rs, ss*} means that we actually have 4 equations for *E*_*mn*_(*t*) and for *I*_*mn*_(*t*), each corresponding to one of the pathogen strains *rr, sr, rs, ss*. In the absence of fungicide treatment, infection occurs with infection rate *β*, with latent rate *γ* (transition from latent infection to symptomatic infection) and removal rate *µ* (transition from symptomatic to removed).

#### Primary infection

The modelled season starts at *T*_*emerge*_ (Table 1), which corresponds to the emergence of ‘leaf five’ (***Elderfield et al., 2018***; ***van den Berg et al., 2013***), rather than the start of the growing season. The dynamics between the start of the growing season and *T*_*emerge*_ are approximated by the initial conditions which are used to subsume both the primary infection and the initial dynamics at the start of the season (***Elderfield et al., 2018***).

The initial infection comes from a primary inoculum *P*_*mn*_ for strain *mn* ∈ {*rr, rs, sr, ss*}. This inoculum is assumed to decay exponentially, at the same rate *v* for all strains. In fact, aside from the effect of the fungicide application, the strains are treated as identical. In particular, this assumes no fitness cost to the presence of fungicide resistance. Letting *t* be the time since the start of the season (measured in degree-days) and *P*_*mn*,0_ be the initial amount of inoculum for strain *mn* ∈ {*rr, rs, sr, ss*}:

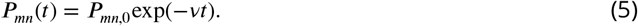

#### Host growth

We define the total amount of tissue

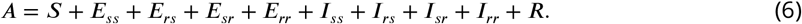

The growth of the wheat crop is given by the following function:

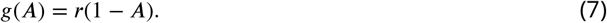

The growth is monomolecular (***Cunniffe and Gilligan, 2010***), and includes density dependence such that the rate of production of host tissue decreases as the total amount of tissue (*A*) increases. We non-dimensionalised the tissue quantities so that the maximum leaf area after growth finishes is 1. This differs from the scale used by ***Hobbelen et al***. (***2013***), where tissue quantities are measured out of 4.2, the maximum leaf area index (due to vertically stacked leaves). The growth function is scaled by a growth rate *r*.

#### Senescence

We use the following senescence function Γ:

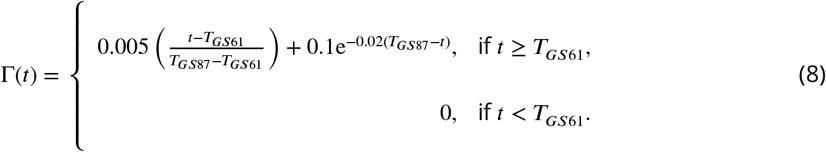

This function is inherited from the models by ***Hobbelen et al***. (***2013***) and ***Elderfield et al***. (***2018***), and represents the rapid increase in the senescence of healthy tissue towards the end of the season. Senescence begins at growth stage 61, *t* = *T*_*GS*61_ (where we use Zadok’s growth scale for the growth of wheat (***Zadoks et al., 1974***)). Senescence is assumed to only affect tissue from the *S* and *E* compartments, but the disease is assumed to disrupt this process meaning that there is no senescence of tissue in the *I* compartment. By harvesting time at *t* = *T*_*GS*87_ we get (almost) complete senescence of the healthy tissue.

### Effect of fungicides

We denote the response of a pathogen strain *m* to the application of a fungicide *F* by *δ*_*F,m*_(*C*_*F*_). The chemical concentration changes depending on the time since the chemical was applied, which means the pathogen response varies with time (Figure 1D). We assume fungicide applications occur instantaneously and that the concentration of fungicide decays exponentially (Figure 1C). Two-treatment fungicide programs were found to balance effective control with resistance management in ***van den Berg et al***. (***2016***), so we focus on strategies which have two spray applications per year. We assume the two fungicide treatments are applied at *T*_*GS*32_ and *T*_*GS*39_ each year (Table 1).

**Figure 1.**
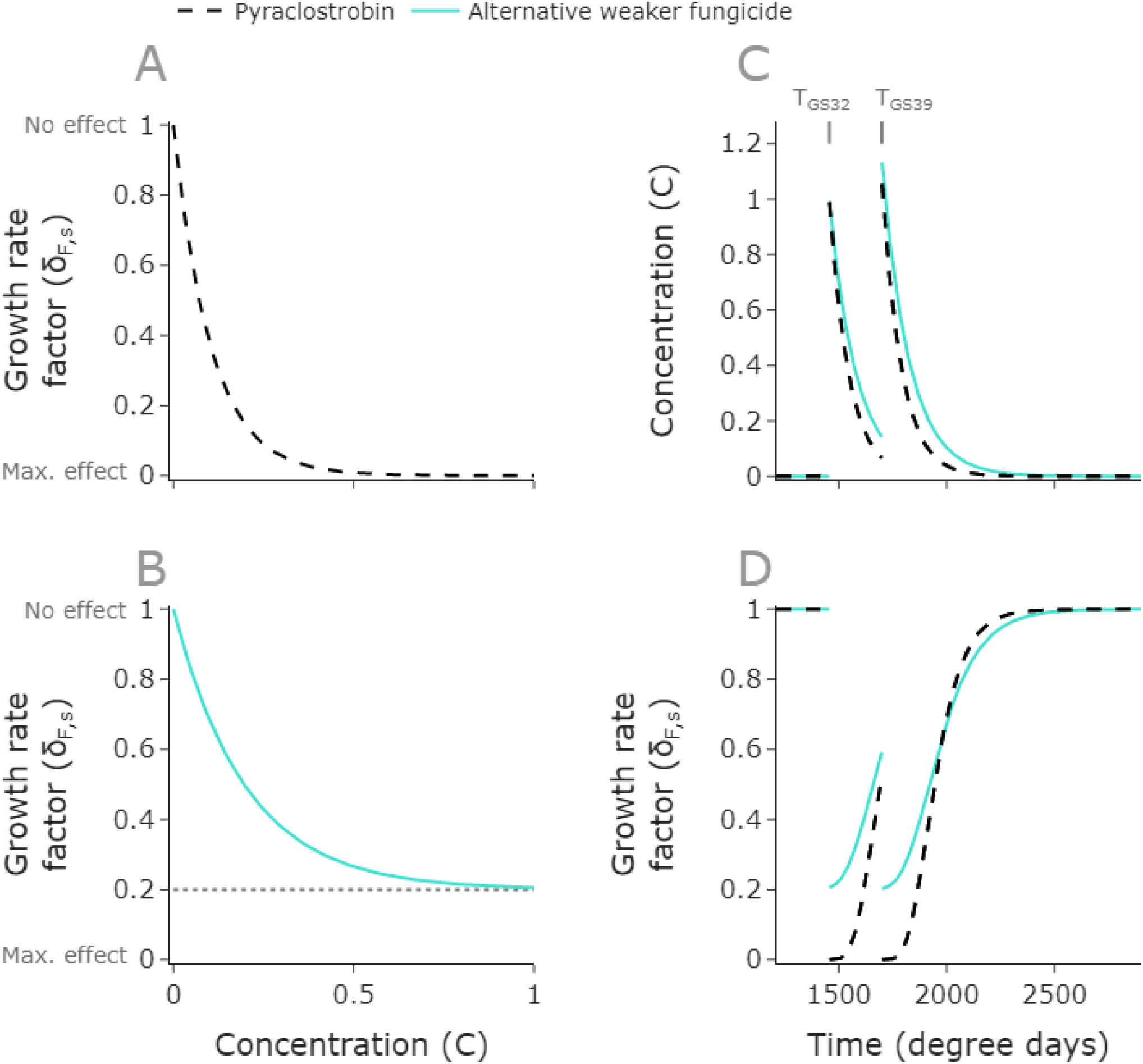
Fungicide applications limit pathogen growth rates before concentrations decay. Dose response curves for pyraclostrobin (**A**) and a weaker but slower decaying alternative fungicide (**B**). We plot *8*_*F, s*_(*C*_*F*_), the multiplier on the growth rate for sensitive strains, so that the growth rate is unchanged (*8*_*F,s*_(*C*_*F*_) = 1) when the chemical concentration is 0, but the growth rate approaches 0 as the concentration of pyraclostrobin increases (**A**). However, the weaker fungicide has a maximum effcacy *w* = 0.8 (**B**), meaning that the growth multiplier approaches 0.2 (dotted line) at high concentrations. The concentration depends on time since application (**C**), starting at 0 at the beginning of the modelled season (1212 degree days as in ***Elderfield et al***. (***2018***)) and increasing instantaneously at *t* = 1456 and *t* = 1700 when the doses are applied, before exponentially decaying with rates that differ here for the two fungicides. The effect of the fungicides correspondingly vary with time since the first spray (**D**). We model scenarios in which two sprays are applied, and here a full dose of both fungicides is applied. **Parameter values**: pyraclostrobin: (*w, θ*, A) = (1, 9.6, 1.11 × 10^−2^); weaker alternative fungicide: (*w, θ*, A) = (0.8, 7, 8 × 10^−3^). Pyraclostrobin parameterisation is as in ***Hobbelen et al***. (***2013***).

The fungicides decrease the rate of the transition of tissue from healthy to exposed (*β*), and from exposed to infected (*γ*), corresponding to both protectant and eradicant activity (Equations 1 - 4). Both transition rates are assumed to decrease by the same amount, as in ***Hobbelen et al***. (***2013***). The fungicide response *δ*_*F,s*_(*C*_*F*_) lies in the interval [0, 1], and multiplies each respective transition rate (Table 2).

**Table 2.**
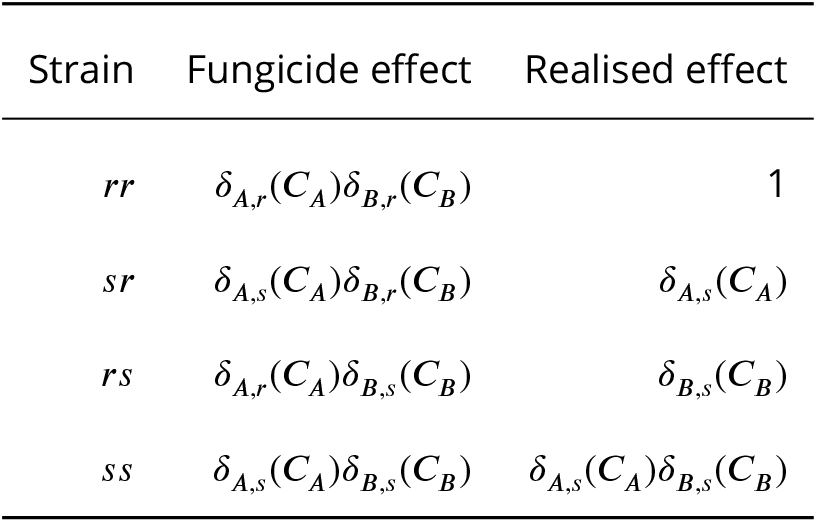
Effect of fungicides on the pathogen strains in the model. Each value in the interval [0, 1] multiplies the rate of the transition of tissue from susceptible to exposed (*β*) and from exposed to infectious (*γ*). This means that the *rr* strain is unaffected by the presence of fungicide (multiplied by 1), whereas the *ss* strain has transition rates reduced by a factor of *δ*_*A,s*_*δ*_*B,s*_ < 1. Note that ‘fungicide effect’ and ‘realised effect’ are equivalent, since *δ*_*A,r*_ = *δ*_*B,r*_ = 1, i.e. resistant strains are assumed to be completely unaffected by an application of fungicide.

For a dose *C*_*F*_ of a fungicide *F*, we use dose responses to sensitive strains (Figure 1A,B) of the type

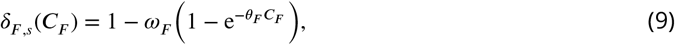

where *δ*_*F,s*_(*C*_*F*_) is the effect on any strains sensitive to it. Here *ω* is the maximum effect of the fungicide and *θ* is a curvature parameter. The curvature parameter *θ* characterises how steeply the curve drops, i.e. how sharply the effect on the pathogen depends on the concentration of chemical (Figure 1A,B). Higher efficacy fungicides have higher values of *ω* and/or *θ*. We predominantly focus on dose responses within a similar parameter range to pyraclostrobin, a high risk strobilurin fungicide modelled in ***Hobbelen et al***. (***2013***). We assume that resistant strains are completely unaffected by an application of fungicide, in the same way as ***Elderfield et al***. (***2018***). This means that *δ*_*F,r*_(*C*_*F*_) = 1 for any fungicide *F* and concentration *C*_*F*_.

The net rate of infection for any strain *mn* is given by the sum of the rates of primary infections and secondary infections. The rate of primary infections is given by *δ*_*A,m*_(*C*_*A*_)*δ*_*B,n*_(*C*_*B*_)*P*_*mn*_(*t*), and the rate of secondary infection is given by *δ*_*A,m*_(*C*_*A*_)*δ*_*B,n*_(*C*_*B*_)*I*_*mn*_(*t*). Initially primary infection contributes highly, but since the primary inoculum decays away exponentially it becomes less important relative to secondary infections as the season progresses.

### Between-season dynamics

Any remaining primary inoculum from the previous winter is assumed to have decayed entirely by the end of the growing season. We assume a constant total initial amount of inoculum in each season (denoted *ψ*_0_), as is used by ***Hobbelen et al***. (***2013***); ***Elderfield et al***. (***2018***).

We keep the proportion of between-season sexual reproduction as a free parameter (*q*_*B*_). Then the remaining proportion 1− *q*_*B*_ of the initial population is assumed to be clonal offspring. Initially we consider *q*_*B*_ = 0 as in ***Hobbelen et al***. (***2013***); ***Elderfield et al***. (***2018***). We later scan over all possible values of *q*_*B*_ to demonstrate the effect of alternative parameter choices.

We denote the proportion of offspring of strain *mn* as *X*_*mn*_, *Y*_*mn*_ for the asexual and sexual cases respectively. The calculation of these quantities is described below. The levels of primary inoculum at the start of the next season are given by:

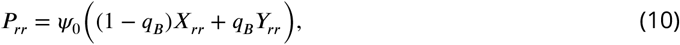

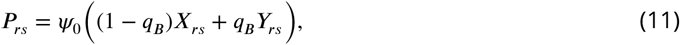

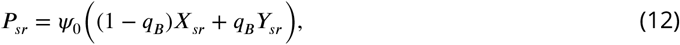

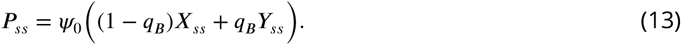

Let 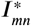 be the level of infection for strain *mn* at the end of the previous modelled season. We assume that the fractions of each pathogen strain at the start of a season are calculated based only on the fractions of infectious tissue infected by each strain at the previous season’s end, as in ***Hobbelen et al. (2011a)***. We also define the sum of all disease strains:

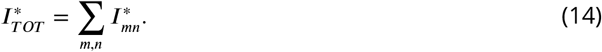

We may also define *F*_*mn*_, the frequency of strain *mn* as a proportion of total disease:

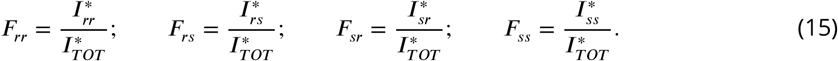

The proportions of asexual offspring, *X*_*mn*_, are simply given by:

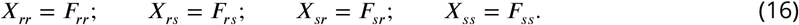

For the sexual offspring *Y*_*mn*_, we assume perfect random mating of unlinked resistance genes in a haploid population, which leads to frequencies as in ***Felsenstein*** (***1965***). The unlinked assumption is valid unless the loci involved are close together on the same chromosome. A full derivation of these frequencies is given in Supplementary Text 1.

Now defining

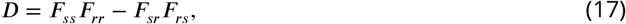

we obtain:

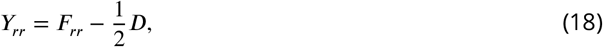

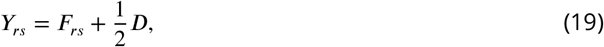

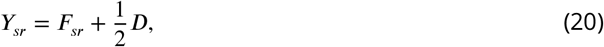

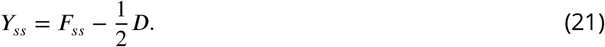

Combining the results for *X*_*mn*_ and *Y*_*mn*_ leads to the following result:

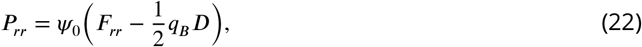

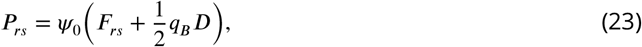

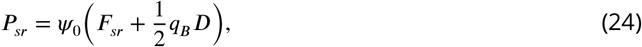

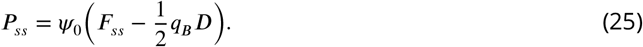

Note that the sum of these values gives *ψ*_0_, the total initial amount of inoculum.

### Measures of strategy performance

#### Effective life

Following ***Hobbelen et al***. (***2013***, 2011a), we use the term *effective life* to denote the number of years for which a fungicide or fungicide application strategy is effective, meaning growers achieve yields above a certain threshold, which we set at 95% of the disease-free yield (***Hobbelen et al., 2013***). We assume that a grower requires a yield above this threshold for economic reasons. We will use effective life to assess the effectiveness of the fungicide mixtures we test.

#### Selection ratio

Again following ***Hobbelen et al. (2011b)***, we use the term *selection ratio* as a measure of how strongly a particular strategy results in selection for the resistant strain. They define the selection ratio in terms of the frequency of the resistant strain before the first spray and at the end of the growing season. We generalise this form to apply to any of our four pathogen strains, so that in any year *N* we use:

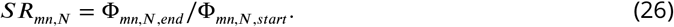

Here Φ_*mn*_ represents the density of strain *mn* as a proportion of total disease, at the start or end of the season. If the selection ratio for a fungicide is greater than 1 then the pathogen strain has increased in frequency in year *N*.

### Example simulation

During the season, susceptible host tissue increases in density initially before natural senescence later on (Figure 2B,C). The four pathogen strains are affected by the fungicide mixture to different extents. The fungicide applications cause the exposed tissue (of sensitive strains) to stop increasing (Figure 2D) and the infectious tissue (of sensitive strains) to decay (Figure 2E). This leads to selection for resistant strains and a gradual loss in yield over successive seasons due to reduced control of the pathogen (Figure 2F). The proportions of the pathogen population carrying resistance to either fungicide increase over successive seasons (Figure 2G). This increase in resistance frequency for either fungicide (e.g. the strains *rs* and *rr* are resistant to fungicide *A*) is approximately logistic and is a result of the selection pressure applied. This is characterised by an initially gradual change in resistance frequency before a rapid increase (Figure 2G).

**Figure 2.**
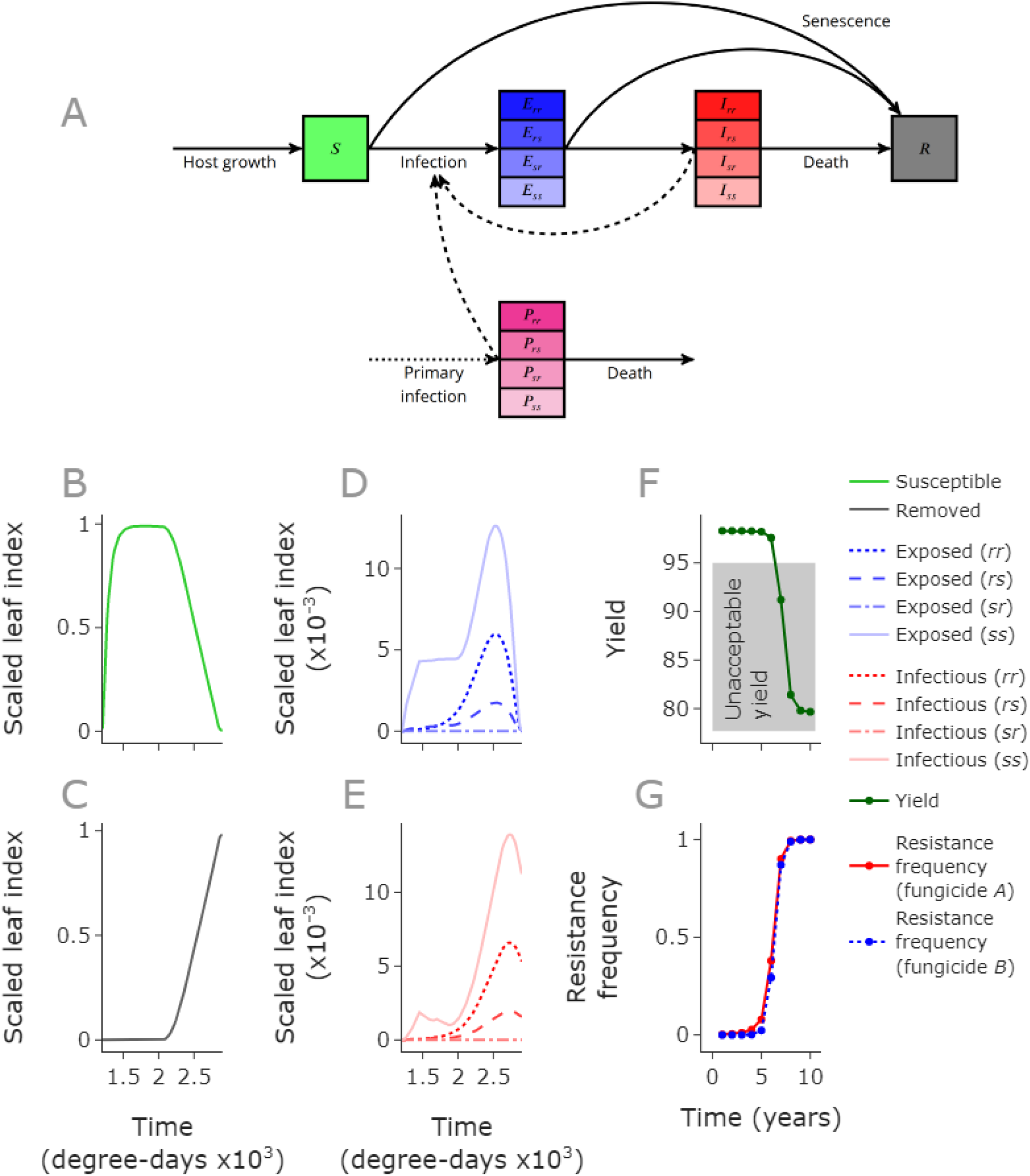
Selection for resistant strains within each season leads to yield losses over time. The diagram (**A**) is a graphical representation of the within-season model. The solid lines represent transitions, the dashed lines effects (for example the amount of infectious tissue affects the rate of infection) and the dotted line represents the instantaneous arrival of primary inoculum (*P*) at the beginning of each season in the model. The pathogen strains are denoted *ss, rs, sr* and *rr*, representing the double sensitive, two single-resistant, and the double resistant strain. *S, E, I, R* represent tissue categories: susceptible; exposed; infectious; and removed (**B**,**C**,**D**,**E**). The modelled season begins at 1212 degree-days (emergence of leaf 5), as in ***Hobbelen et al***. (***2013***), with fungicide applications at *t* = 1456 and *t* = 1700. Over successive seasons, the yield declines to below the ‘unacceptable yield’ threshold (95%, **F**) as the resistance frequencies increase (**G**). The resistance frequencies for a fungicide are the proportion of total disease caused by strains resistant to that fungicide (e.g. for fungicide *A* these are strains *rs* and *rr*). **Parameter values**: default values as used by ***Hobbelen et al***. (***2013***), doses: (1, 0.5). Fungicides *A* and *B* parameterised to match efficacy of pyraclostrobin. Initial resistance frequencies: (*rr, rs, sr*) = (10^−8^, 10^−3^, 10^−6^) – note that these are slightly higher than the default values used in later figures. The disease progress curves are for the 5th growing season. Choosing higher resistance frequencies and a later growing season means the resistant strains are sufficiently high in density to be visible on the same scale as the sensitive strain.

## Results

### Finding an optimal high-risk mixture strategy

We seek the optimal dose-pairing to minimise selection for resistant strains and prolong the effectiveness of the mixture while providing sufficient disease control to be economic. We restrict our search to fixed-dose strategies; those strategies for which each year the same dose choices are used, although the dose of fungicide *A* and of fungicide *B* may differ to each other in the mixture.

Previous work on high-risk fungicide mixtures has neglected the scenario where frequencies of the single resistant strains differ initially. We seek to determine the optimal strategy depending on these initial conditions, as well as fungicide parameterisation (dose response and decay rate) and the balance of asexual and sexual pathogen reproduction. Initially we assume that there is no sexual reproduction in the pathogen population, but we relax this assumption later.

#### Dose space

We define ‘dose space’ as the set of pairs of fungicide doses which take values between 0 and 1, since we assume a “full dose” of 1 is the maximum dose that would be permitted (Figure 3A,B,C). This allows us to describe all permissible dose-pairs for our fixed fungicide strategy. Different choices lead to different effective lives (the number of years for which this strategy gives acceptable yields). We seek the optimal region within dose space, which is the region containing dose combinations which lead to the longest effective life for the mixture. This is a region rather than a single pair of doses, since there may be multiple dose combinations which break down in the same (optimal) year (Figures 3A, 4A-D).

**Figure 3.**
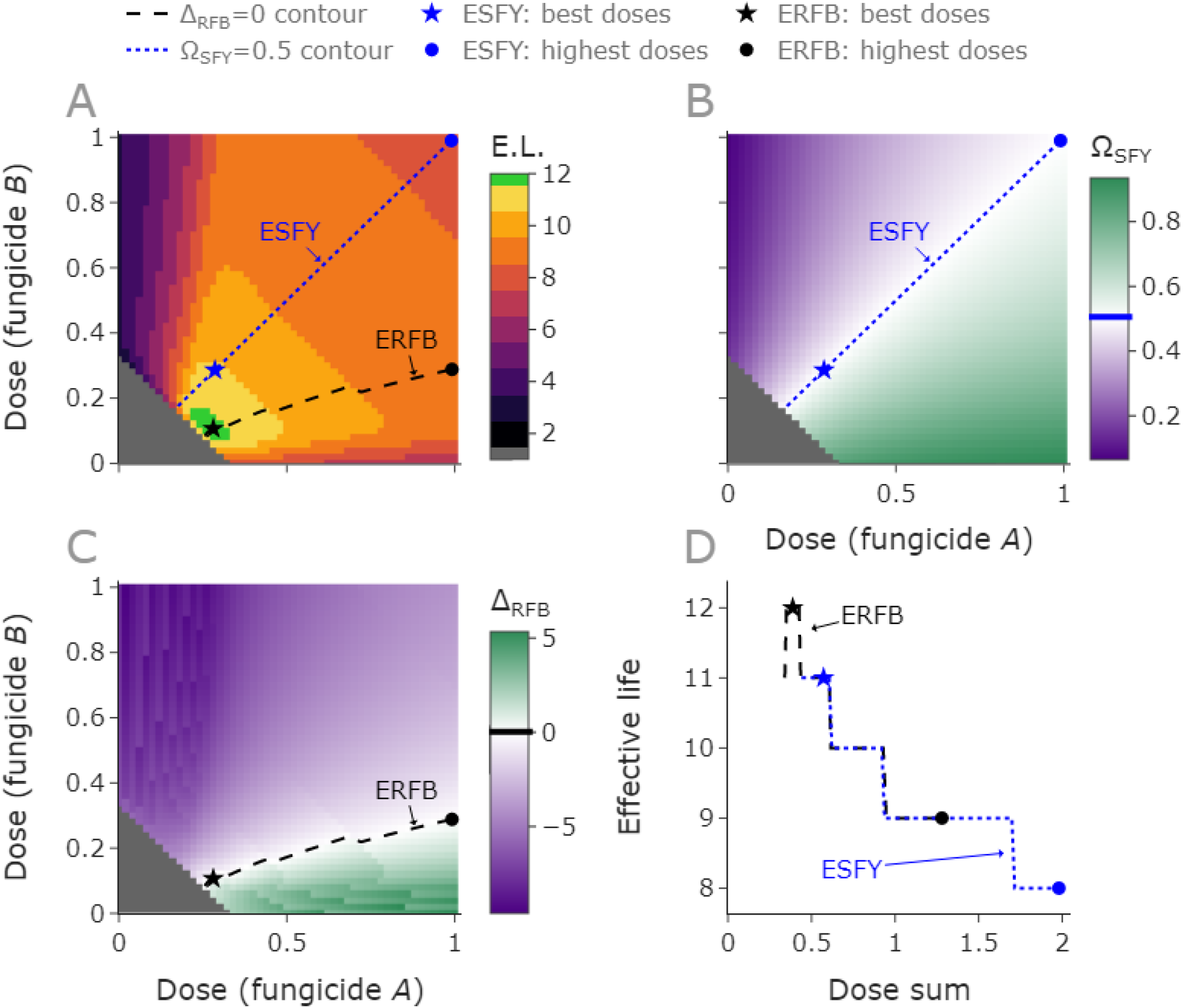
The equal single resistant frequencies at breakdown strategy is optimal in some cases where equal selection in first year is sub-optimal. **A, B, C**: ‘Dose space’ for the scenario where the two fungicides are assumed to act with equal effcacy, but the initial levels of resistance are unequal, with greater levels of resistance to fungicide *B* initially. The grey region is doses for which the mixture isn’t suffciently strong to give acceptable yields even in the first year. The optimal region is the green region with the highest effective life of 12 years (**A**). Examples of the best doses along each contour are denoted by stars, and the highest permissible doses (strongest mixture) are denoted by circles. The dotted blue line represents doses where selection is equal after the first year of treatment (the ESFY strategy). There are no doses along this contour which lie in the optimal region of dose space. This is an example of a case in which the recommendation given by ***Hobbelen et al***. (***2013***) fails. The values of Ω_*SFY*_ in dose space are shown in (**B**). The dashed black line represents those doses which lead to equal resistance frequencies for the two fungicides in the breakdown year (the ERFB strategy). Note that there are doses along this contour which fall in the optimal region (**A**). The Δ _*RFB*_ contour in dose space is shown in (**C**). This strategy reacts much more to differing initial levels of resistance. The effective lives are shown against different values of ‘dose sum’ which is the sum of the doses of fungicide *A* and *B*, as we move along the Ω_*SFY*_ = 0 and Δ _*RFB*_ = 0 contours (**D**). **Parameter values**: both fungicides with dose response curves and decay rate parameter as per pyraclostrobin (***Hobbelen et al., 2013***): (*θ, w*, Λ) = (9.6, 1, 0.0111), initial resistance frequencies: (*I*_*rs*_, *I*_*sr*_, *I*_*rr*_) = (10^−7^, 10^−3^, 1.0010009487970706 × 10^−10^). ***Hobbelen et al***. (***2013***) use default initial values of 10^−5^ for the single resistant strains - we chose values 10^2^ bigger/smaller than these values.

**Figure 4.**
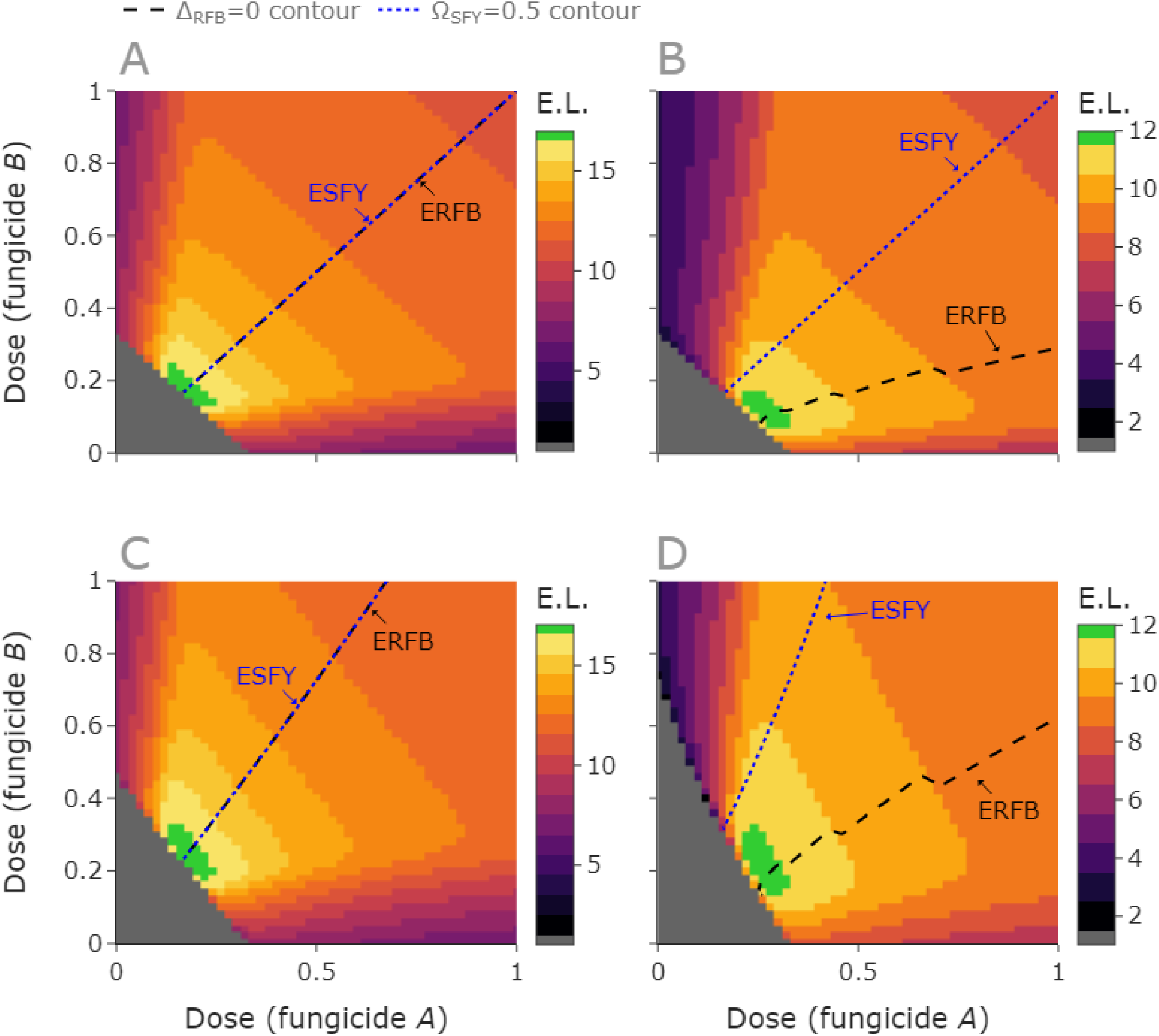
When does equal resistance at breakdown outperform equal selection in the first year? When the fungicides have identical dose responses, and the initial (single) resistance frequencies are the same, the ERFB and ESFY strategies are equivalent, and both are optimal (**A**). When the initial frequencies differ, but the fungicide parameterisations are the same, the strategies differ (**B**), as demonstrated by the difference in location of the ERFB and ESFY contours. Here ERFB outperforms ESFY; the ERFB contour passes through the optimal region but the ESFY contour does not. When the fungicide parameterisations differ, but initial (single) resistance frequencies are the same, the ERFB and ESFY strategies are very closely aligned, and both optimal (**C**). When both fungicide parameters and initial (single) resistance frequencies differ, again the strategies differ, but ERFB outperforms ESFY (**D**). **Parameter values**: **A**,**B**: Both fungicides with dose response curves and decay rate parameter as per pyraclostrobin (***Hobbelen et al., 2013***): (*θ, w*, Λ) = (9.6, 1, 0.0111). **C**: fungicide A as per pyraclostrobin, fungicide B: (*θ, w*, Λ) = (7.5, 0.95, 0.0111). **D**: fungicide A as per pyraclostrobin, fungicide B: (*θ, w*, Λ) = (7, 0.8, 0.0111). **A**,**C**: Equal initial single RFs; (*I*_*rs*_, *I*_*sr*_, *I*_*rr*_) = (10^−7^, 10^−7^, 9.992007221626409 × 10^−15^). **B**,**D**: Different initial single RFs; (*I*_*rs*_, *I*_*sr*_, *I*_*rr*_) = (10^−7^, 10^−3^, 1.0010009487970706 × 10^−10^).

The position and size of the optimal region differs for different initial resistance frequencies, different fungicide parameters or different proportions of between-season sexual reproduction (for example the optimal regions in Figure 4A-D all differ). To find the optimal region, we use a grid of dose choices which discretises both concentrations and considers pairs of doses of each chemical (labelled fungicide *A* and *B* respectively). This ‘brute-force’ method allows us to simply read off which dose combinations are best for a given model parameterisation. This method was previously used by ***Hobbelen et al***. (***2013***), although we use a finer grid of 51 as opposed to 11 (i.e. for each parameterisation we consider 51^2^ = 2601 rather than 11^2^ = 121 pairs of doses). We also seek a more biologically-motivated and general method to characterise the optimal dose region, depending on characterisations of the pathogen and fungicides (initial levels of resistance, mode of reproduction, decay rates and dose-response curves).

#### Strategies to test

We will compare the following candidate strategies:

- Equal selection in the first year (ESFY),
- Equal resistance frequencies at breakdown (ERFB).

We aim to see how these strategies perform relative to the grid search.

#### Equal selection in the first year

When strains are completely resistant to fungicide *A* and *B*, ***Hobbelen et al***. (***2013***) assert that the most durable strategy exerts an approximately equal selection pressure on both single-resistant strains. They considered situations where the frequencies of the single resistant strains were equal initially and show how equal selection in the first growing season gives the best outcome. Therefore we choose this as our first candidate strategy.

To explore this strategy we define a metric Ω_*SFY*_ :

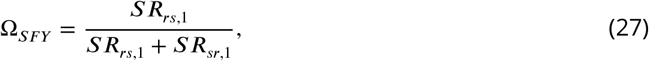

where *SR*_*mn*,1_ is the selection ratio for strain *mn* in year 1, as defined in Equation 26. Then Ω_*SFY*_ = 0.5 means there is equal selection for both single resistant strains in the first year of chemical application (Figure 3B), whereas Ω_*SFY*_ > 0.5 means there is greater selection for fungicide *A*, and Ω_*SFY*_ < 0.5 means there is greater selection for fungicide *B*. There is a curve in dose space defined by the doses satisfying Ω_*SFY*_ = 0.5. In the equal dose-response, equal initial resistant frequency case, this contour is the line *y* = *x*, for sufficiently large pairs of doses that we get an acceptable yield. If the initial resistance frequencies are low enough that density dependent effects have a negligible effect on selection, the line still falls approximately along *x* = *y* even if the frequencies differ (Figure 3).

Minimal doses (that still achieve sufficient disease control) along the Ω_*SFY*_ = 0.5 contour are equivalent to what was recommended previously (***Hobbelen et al., 2013***). We will explore this strategy (equal selection in first year, low dose), and generalise it slightly to additionally explore a strategy which takes any dose along this contour rather than only considering minimal doses (equal selection in first year). Considering all doses on the contour gives an increased chance of finding good dose combinations, and we are also interested in the relative performance of higher dose mixtures relative to lower dose mixtures. We define ‘low doses’ by considering the minimum viable and maximum permitted so-called ‘dose sum’ along this contour (we define dose sum as dose of fungicide *A* + dose of fungicide *B*). Low doses are within the lowest third (arbitrarily chosen) of dose sums which give acceptable control, a range which depends on the parameterisation.

If the initial frequencies of the resistant strains differ – for example for pairs of fungicides that were introduced to market at different times – the ESFY recommendation sometimes fails to give the optimal outcome (Figures 3A, 4B,D).

#### Equal resistance frequencies at breakdown

We propose an alternative strategy – one that ensures resistance frequencies are equal in the first year where the yield becomes unacceptable (the ‘breakdown year’). This takes into account the effect of the strategy over its entire course rather than only its first year, and it more effectively accounts for differing initial levels of resistance to the two fungicides. This strategy performs as well or better when compared to the ESFY strategy, using the default parameterisation and a range of initial conditions (Figure 3A,C,D).

To explore this strategy, we will define another quantity: Δ_*RFB*_. This is defined in terms of the difference between the (logits of the) single resistance frequencies at breakdown:

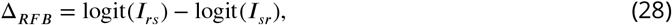

where

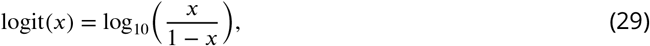

and *I*_*rs*_, *I*_*sr*_ are the densities of single resistant strains to fungicides *A* and *B* respectively, measured at the end of the breakdown year. This quantity informs us about the state of the system in the breakdown year, and whether our strategy led to a greater degree of resistance to one fungicide compared to its mixing partner. The quantities Ω_*SFY*_ and Δ_*RFB*_ are conceptually similar, focusing on the effect of the strategy on the single resistant strains. However Δ_*RFB*_ relates to *resistance frequency* in the *breakdown* year as opposed to *selection* in the *first* year (Ω_*SFY*_). We use a different function involving logit(*I*_*mn*_) for Δ_*RFB*_ because the densities of the single resistant strains can be as small as 10^−10^, so a logit scale is more appropriate, whereas because the selection is calculated in terms of a ratio, it typically lies between 1 and 10. We choose a logit scale rather than logarithmic because the growth of each single resistant strain is approximately logistic.

If the resistant frequencies are equal in the breakdown year then Δ_*RFB*_ = 0. If there is more resistance to fungicide *A* than *B*, Δ_*RFB*_ is positive and if there is more resistance to fungicide *B* than *A* then Δ_*RFB*_ is negative. There is a contour in dose space described by Δ_*RFB*_ = 0 (Figure 3A,C). We will refer to this as the Δ_*RFB*_ contour.

For the identical fungicide pair model parameterisation, some points along the Δ_*RFB*_ contour lie in the optimal region (Figure 3A). For these points, resistance frequencies in the breakdown year are equal and our strategy is optimal. Note that there are also points along the Δ_*RFB*_ contour that are not optimal – this is because the strength of the mixture must be carefully chosen according to the fungicide/pathogen parameters. This process is examined in the wider parameter scan below. Note that in Figure 3D the optimal mixture strength is slightly higher than the weakest possible mixture strength. This is because the weakest possible mixture strength gives a yield in the first year of 95%. Any small increase in resistance then causes strategy failure. A slightly higher mixture strength achieves sufficiently good control to withstand these small changes in resistance in the early years, but much higher mixture strengths lead to stronger selection for resistant strains and hence much poorer resistance management. This trade-off causes the non-monotonic shape of the ERFB line in Figure 3D, which shows how effective life varies with mixture strength (parameterised as dose sum). Although ***Hobbelen et al***. (***2013***) suggest that the optimal choice of dose is just high enough to provide effective control, this result suggests doses slightly higher than minimal are optimal. This result may have been obscured by the coarser 11 by 11 grid in dose space in that work.

The Δ_*RFB*_ contour is smooth within each effective life region (Figure 3A), but jumps as it moves into regions with a different effective life (so different breakdown year). This is because the values of Δ_*RFB*_ are found using the relative amounts of single resistant strains in the breakdown year; in each different region the breakdown year differs and the relative amounts do not align exactly in different seasons.

If initial resistance frequencies and fungicide parameterisations are the same, this recommendation is equivalent to equal selection in the first year as found in ***Hobbelen et al***. (***2013***) (Figure 4A). However, the ESFY recommendation may not work if the initial resistance frequencies and fungicide parameterisations are not identical. If the initial resistance levels differ then ERFB may outperform ESFY (Figure 4B,D). However, we note that if the fungicide parameterisations differ but the initial resistance levels are the same, then the two strategies are very closely aligned (Figure 4C). This is because if selection is equal every year then resistance frequencies will be equal at breakdown. Density dependent effects mean that equal selection in the first year does not guarantee equal selection in subsequent years, but this effect is small enough to be ignored in many cases. Equalising resistance frequencies at breakdown avoids the situation where resistance develops much faster to one fungicide in the mixture, causing it to become ineffective. That would lead to a loss of control for that fungicide and a lack of protection offered to its mixing partner, causing the mixture to fail more quickly than if resistance frequencies are equal at breakdown.

### The effect of sexual reproduction

We explored the effect of sexual reproduction for a range of initial frequencies and fungicide parameterisations, which were selected randomly and independently (Table 3). We tested the maximum effective life from a grid of 21 by 21 doses with 11 different values of *q*_*B*_ between 0 and 1 for each set of parameters. Three such examples which demonstrate the possible behaviours are shown in Figure 5. We refer to each example as a ‘replicate’.

**Table 3.** Ranges/values taken by parameters in the scan over values of the between-season sexual reproduction proportion. The range of fungicide decay rates depends on Λ_0_, set as 1.11 × 10^−2^ degree days^-1^ as in Table 1. The value of the double resistant strain is fixed such that = 0 initially, and then we explore what happens when the double resistant strain is at a density much lower or higher (10^−5^ or 10^5^ times lower/higher) in each case (see Figure 5). Parameters were independently generated and parameter sets were accepted as long as the yield in the first year at full dose was at least 95% of the disease free yield, so that there was a valid strategy possible.

| Parameter | Symbol | Range/Value | Distribution |
| --- | --- | --- | --- |
| Pathogen between-season sex proportion | $q_B$ | [0, 1] | Uniform |
| Single resistant strain initial frequencies | $I_{mn}(0)$ | $[10^{-8}, 10^{-4}]$ | Log-uniform |
| Double resistant strain initial frequencies | $I_{rr}(0)$ | Varied | Fixed by $sr$, $rs$ strains |
| Fungicide decay rates | $\Lambda_i$ | $\left[\frac{1}{3}\Lambda_0, 3\Lambda_0\right]$ | Uniform |
| Fungicide asymptotes | $\omega_i$ | [0.4, 1] | Uniform |
| Fungicide curvatures | $\theta_i$ | [4, 12] | - |

**Figure 5.**
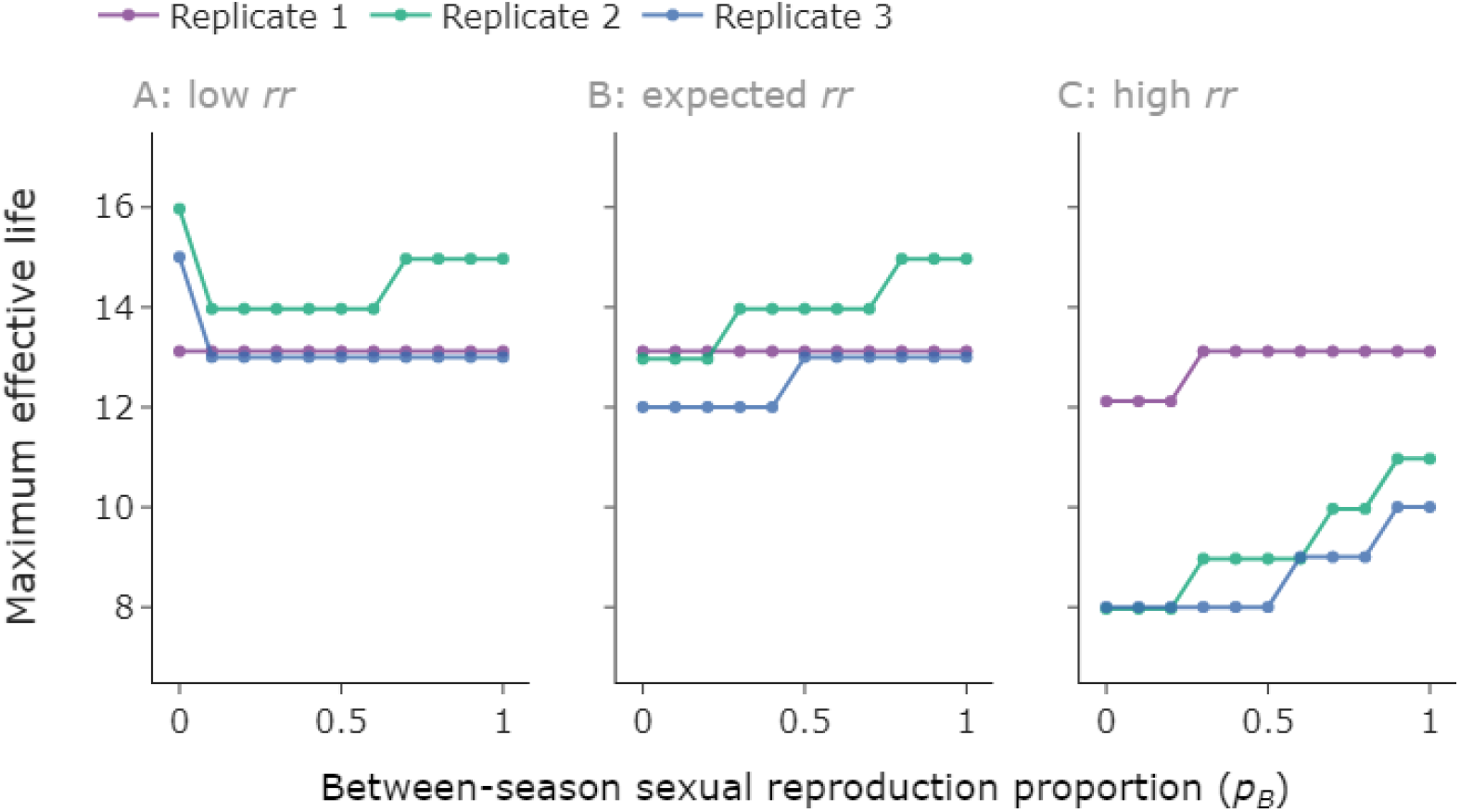
Effect of sexual reproduction proportion depends on the initial level of the double resistant strain. We examine three randomly generated replicates (Table 3). For each replicate, we consider three possibilities: low, expected and high double resistant density (**A**/**B**/**C**). To avoid the lines overlapping we shifted each replicate vertically by a small amount, but each effective life should be an integer. **Parameter values:** random fungicide parameter values and single resistant frequencies, generated from distributions described in Table 3. **A**: initial double resistant strain (*rr*) density 10^−4^ times lower than value from linkage equilibrium (VLE); **B**: initial *rr* density at VLE; **C**: initial *rr* density 10^4^ times higher than VLE. Values: (*p*_*rr*_, *p*_*rs*_, *p*_*sr*_, Λ_*Λ*_, A_*B*_, *w*_*A*_, *w*_*B*_, *θ*_*A*_, *θ*_*B*_) = **Replicate 1**: (7.049916 × [10^−19^ 10^−15^ 10^−11^], 5.545997 × 10^−7^, 1.269711 × 10^−8^, 0.009758, 0.022030, 0.729797, 0.661193, 7.362942, 6.642679); **Replicate 2**: (1.085892 × [10^−15^ 10^−11^ 10^−7^], 1.596583 × 10^−6^, 6.801290 × 10^−6^, 0.007417, 0.009834, 0.574543, 0.706497, 11.143576, 11.170345); **Replicate 3**: (2.445955 × [10^−15^ 10^−11^ 10^−7^], 7.155682 × 10^−5^, 3.417953 × 10^−7^, 0.005803, 0.006279, 0.875035, 0.717337, 8.544356, 11.404773).

Initially we explored the initial value of the double resistant strain we expect at linkage equilibrium; meaning the initial value satisfied *I*_*rr*_*I*_*ss*_ = *I*_*sr*_*I*_*rs*_. These are the frequencies we would find in the absence of selection and if frequencies were combined at random. We also demonstrate the effect of increasing *q*_*B*_ when *I*_*rr*_ was not at linkage equilibrium, which could occur due to stochasticity or due to selection within a season, in the scenario where between-season reproduction was not entirely sexual. We tested two alternative scenarios: (i) low double resistant initial density (ii) high double resistant initial density. These were set to be 10^−4^ times lower/10^4^ times higher than linkage equilibrium respectively. Although these are exaggerated values compared to the variation from linkage equilibrium which might be expected in practice, they demonstrate the way the between-season reproduction influences the model output (Figure 5).

When the initial proportion of double resistant was greater than the value expected from linkage equilibrium, higher proportions of sexual reproduction slowed the increase in the double resistant strain (Figure 5C). The maximum effective life often also increased with higher proportions of sexual reproduction when the initial proportion of double resistant was equal to the linkage equilibrium value (replicates 2,3 Figure 5B). This increase in effective life is because the sexual reproduction step reduces the amount of the double resistant if there is already an increased proportion of double resistant strain relative to single resistant strains (see Supplementary Text 2). This is clear from the equations for between-season reproduction:

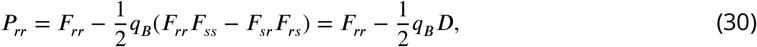

which shows that if *D* = *F*_*rr*_*F*_*ss*_ − *F*_*sr*_*F*_*rs*_ > 0, then the sexual reproduction step will suppress the double resistant strain.

For replicate 1 (Figure 5B), there is no effect on effective life as *q*_*B*_ varies (Figure 5A,B), because the suppression of the double resistant strain is offset by an increase in one or both of the single resistant strains (Supplementary Text 2). Whether the single resistant strains are sufficient to lead to strategy failure depends on whether the fungicides in the mixture are low efficacy, so that each mixing partner cannot adequately control the single resistant strain which is sensitive to it.

For low initial incidences of the double resistant (*rr*) strain, between-season sexual reproduction sometimes acts to decrease the effective life (e.g. replicates 2 and 3, Figure 5A). Interestingly, in replicate 2 there is a non-monotonic response (Figure 5A). This is because any amount of sexual reproduction greatly increases the level of the double resistant strain after the first between-season sexual reproduction step. After this, higher levels of sexual reproduction tend can slow the increase in the double resistant strain that was caused by selection, or it can have no effect, as described above. This behaviour is explored in Supplementary Text 2. As the level of double resistant strain increases, the effective life observed tends to decrease. For example in replicate 2 it decreases from 16 (Figure 5A) to 8 (Figure 5C) when *q*_*B*_ = 0.

Previously mixtures were found to outperform alternations in the absence of sexual reproduction (***Elderfield et al., 2018***; ***Hobbelen et al., 2013***). When between-season reproduction is included, mixtures still outperformed alternations in every case across 100 tested randomly generated scenarios (Supplementary Text 3).

### Generalising to different pathogen and fungicide parameterisations

We tested the robustness of the observation that there are optimal dose combinations along the Δ_*RFB*_ contour, and that the ERFB strategy outperforms the ESFY, by testing a wide range of possible parameter values. We performed a randomisation scan test across different pathogen and fungicide parameter values to check whether this recommendation applies in different scenarios, including varying levels of between-season sexual reproduction, initial levels of resistance, and fungicide dose-responses and decay rates (Table 4). We have seen that the introduction of between-season reproduction can have an effect on the maximum effective life (Figure 5), and seek to show that the ERFB strategy is still optimal despite this effect and is robust to different initial conditions and fungicide parameter values.

**Table 4.**
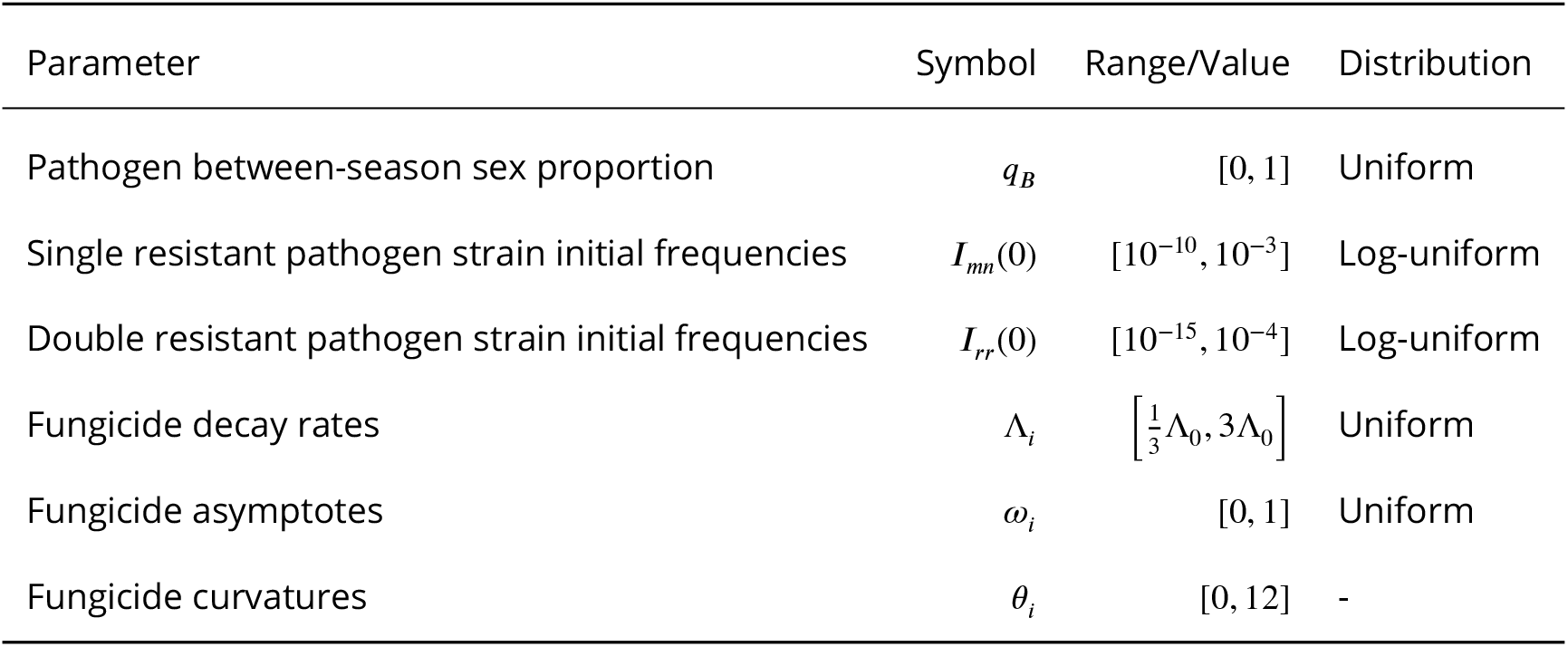
This table shows the ranges/values taken by parameters in the ERFB robustness scan. Table 1 describes the other default parameter values, their sources and their units. The range of fungicide decay rates depends on Λ_0_, set as 1.11 × 10^−2^ degree days^-1^ as in Table 1. Certain parameter combinations were excluded to ensure that both ERFB and ESFY were possible, as described in the main text.

#### Choice of pathogen parameters

The model uses a proportion of between-season sexual reproduction *q*_*B*_. In this section we allow *q*_*B*_ to vary between 0 and 1, since there is no agreed-upon experimental value that we could use. To characterise the pathogen population with respect to the two fungicides, we allow the initial level of resistant strains to also vary. We independently choose values for the proportion of the population taken up by the two single resistant strains and for the double resistant strain. The remaining proportion of the population is the double sensitive strain.

#### Choice of fungicide parameters

The fungicides in the model are characterised by three parameters: the decay rate of the chemical; the curvature parameter; and the asymptote of the dose-response curve. Fungicides with a higher curvature or asymptote value are more efficacious. Longer decay rates also increase the efficacy of a fungicide. We vary all three parameters for each fungicide in the scan.

#### Parameter scan process

For each simulation in the ensemble, we sampled randomly from independent distributions for each parameter (Table 4). We repeated 500 times, giving 500 scenarios with different fungicide and pathogen characteristics as described above. In each case, we compared the best effective life from the grid to the best ERFB and ESFY outcomes.

For each parameter combination, we ran the model over a 51 × 51 grid of dose pairings. We also found the ERFB and ESFY contours in dose space, and compared the optimal effective life from doses on these contours to the optimal value from the grid. The ERFB and ESFY contours are not constrained to lie on the set of pairs of doses we used for our grid, and using only these values would not necessarily lead to a smooth curve in dose space. Instead we characterised points on each contour in terms of the total dose applied, which approximately relates to the strength of the mixture. In particular, we identified a set of lines in dose space along which *C*_*B*_ = *C*_*DS*_ − *C*_*A*_, where *C*_*A*_, *C*_*B*_ are the concentrations of fungicide *A* and *B*, and *C*_*DS*_ is the so-called ‘dose sum’ (i.e. dose of fungicide *A* + dose of fungicide *B*). For each such line we used a numerical optimiser to find the single pair of doses which minimised the squared deviation from the relevant value (0 or 0.5 for Δ_*RFB*_ and Ω_*SFY*_ respectively). The optimiser used was ‘minimize’ from the open-source Python package ‘scipy.optimize’.

This process found this point precisely rather than to the ±0.02 that would be obtained using the grid. Repeating the process for a total of 100 dose sums *C*_*DS*_ identified a smoothly varying contour of pairs of doses precisely along the contour and avoiding the rounding error that would have been associated with ‘snapping’ to the grid of points. This meant that when effective lifetimes were compared, it was possible for the tested strategies to find a higher, equal or lower optimal value compared to the grid.

#### Restriction on parameter space

We restricted the parameter scan to only those fungicides for which equal selection in first year and equal resistance at breakdown was possible. For certain particularly weak fungicides, the minimum dose of the mixing partner (required to attain acceptable yield) was sufficiently large that equal selection was not possible, or if initial levels of resistance to the mixing partner were high then the mixing partner would always break down first.

We excluded any of these problematic parameter combinations by restricting our scan to those for which a full dose of either chemical on its own could give sufficient control (> 95%) at least in the first year of application. This restriction corresponded to excluding fungicides which were not strong enough to provide adequate control if used as a solo product in the first year, and meant that the corners (1,0) and (0,1) of dose space correspond to doses that give acceptable control. The restriction was sufficient to ensure that both strategies were viable in each scenario tested.

#### Parameter scan results

Here we present the results of the parameter scan demonstrating how often the ERFB and ESFY strategies are optimal for randomly selected parameter values, rather than only showing ERFB worked when using the default parameter values. We want to establish which parameters affect the benefit to using ERFB over ESFY, and whether there are any cases where ERFB fails to be optimal. We find that ERFB is almost always optimal and explore the few cases where it is sub-optimal in Supplementary Text 5.

The ERFB strategy is optimal in 99% of cases (i.e. 495 out of 500). This means there is at least one point along the Δ_*RFB*_ contour which gives an effective life at least as good as the optimum effective life from the grid. The ESFY strategy is less effective, performing optimally in 72% of cases. These results are summarised in Table 5. There were only 5 cases where ERFB was sub-optimal (by one year only). These cases are described in Supplementary Text 5. Although there are many cases where the ESFY strategy performs worse than the ERFB contour, it is usually only one year away from the optimum, and is at most 5 years from the optimum. There is a single case in which the ESFY strategy outperformed the ERFB strategy by one year. This case is described in Supplementary Text 5.

**Table 5.**
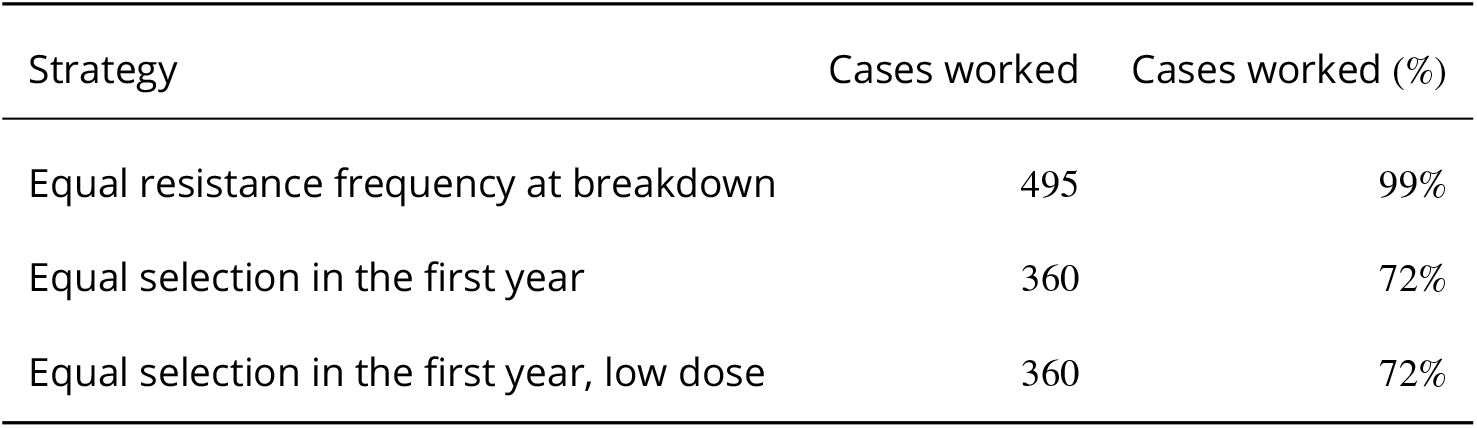
This table shows the results of the ERFB robustness parameter scan, which tested 500 different randomly generated scenarios. ‘Cases worked’ is the number/percentage of runs in which the strategy was at least as durable as the optimal strategy from the 51 × 51 grid. ERFB worked in all but 5 exceptional cases which were within one year of the optimum, whereas ESFY (low dose) and ESFY (all doses) both failed to give optimal results in 28% of cases. ERFB worked every time to within one year whereas both ESFY tactics were at least two years from the optimum in over 10% of cases (12.4% for all doses, 12.6% for low doses only). The cases where ERFB was sub-optimal are addressed in Supplementary Text 5.

There was a relationship between the difference in initial frequencies (on a log scale) and improvement in performance from using ERFB instead of ESFY (Figure 6A). This is as expected -– the strategy which takes differences in initial frequencies into account (ERFB) more effectively performs better relative to the other strategy (ESFY) when these differences are larger. There is less of a clear pattern when looking at the initial values of the double resistant strain, although notably for very large initial values (greater than 10^−5^) there is no difference in strategy performance, since the double resistant strain dominates and causes the loss of control (Figure 6B). This is because the improvement of ERFB on ESFY is usually caused by changes in the relative amounts of fungicide *A* and *B* in the mixture (due to the different positions of the contours). Both fungicide *A* and *B* are ineffective against the double resistant strain, so changes in the relative amounts of each do not increase the effective life if the double resistant strain is the dominant cause of loss of control. For smaller levels of the double resistant frequency, larger differences in performance are sometimes observed. There is no clear pattern in the performance when looking at differences in the fungicide asymptote parameter (Figure 6C), which again suggests that the initial frequencies are the most important feature in terms of whether ERFB outperforms ESFY.

**Figure 6.**
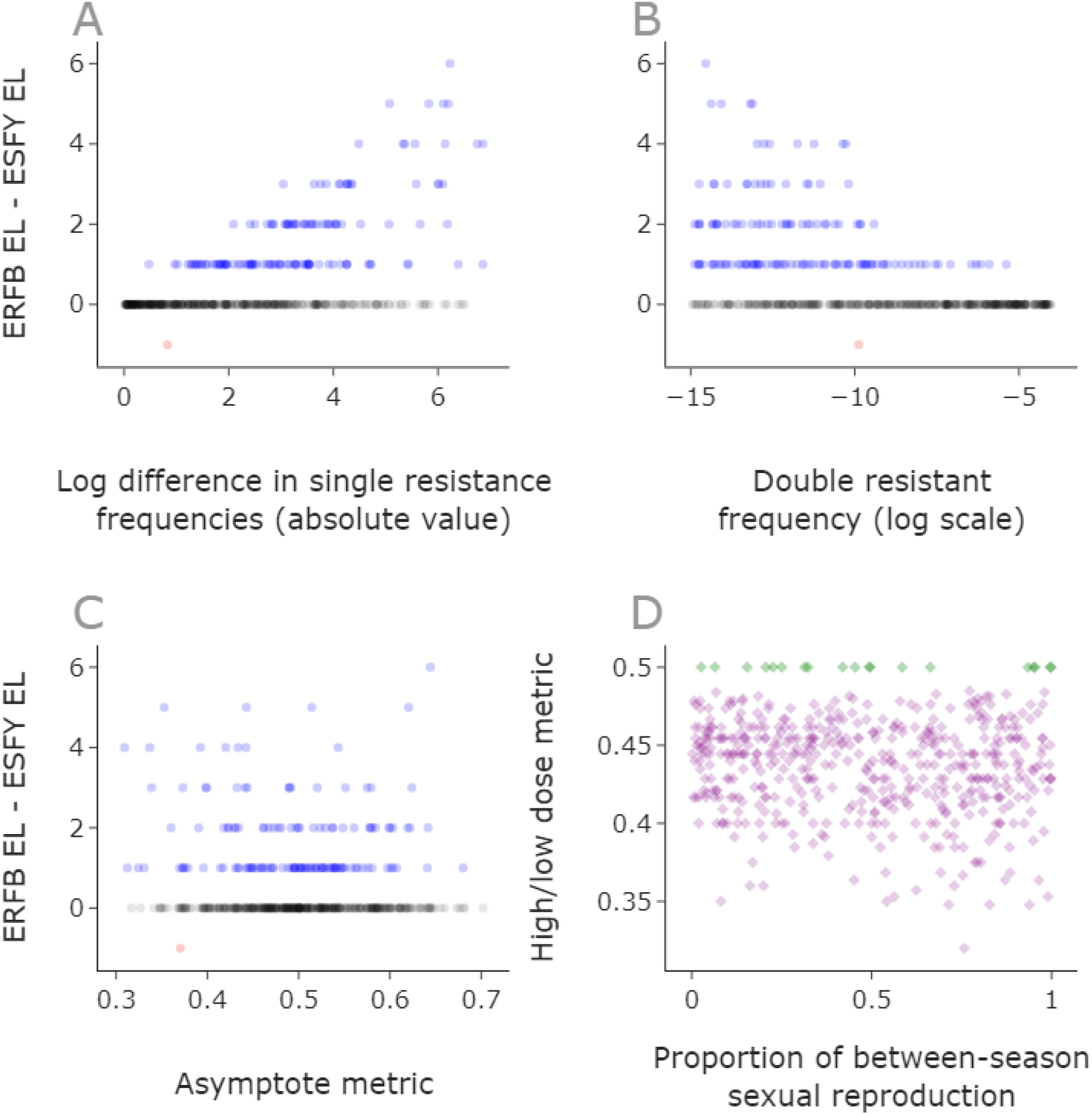
Initial resistance frequencies drive the improvement in the ERFB strategy over ESFY. **A**,**B**,**C:** we plot the difference between the effective life of the ERFB strategy and the ESFY strategy for each of the 500 scenarios in the parameter scan (Table 4). The equal resistance at breakdown strategy always produces an equal (black) or better (blue) effective life when compared to equal selection in first year (all doses) strategy. The quantity on the *x*-axis of subplot **A** is the log difference in the single resistance frequencies |log(*I*_*rs*,0_) − log(*I*_*sr*,0_)|. Subplot **B** shows the double resistant frequency on the *x*-axis, while subplot **C** shows the ‘asymptote metric’, *w*_*A*_/(*w*_*A*_ + *w*_*B*_), where *w*_*F*_ is the asymptote of the dose-response curve for a fungicide *F* (Table 1/Figure 1). Values away from 0.5 indicate greater differences in maximum effcacy for the two fungicides. There is a single case where ESFY outperforms ERFB (red), so that ERFB EL - ESFY EL is negative – this case is explored in Supplementary Text 5. In the examples tested in the parameter scan, low doses were equal (green) or better (purple) than high doses (**D**). The quantity on the *y*-axis of subplot **D**, ‘high/low dose metric’ is *EL*_*low*_/(*EL*_*low*_ + *EL*_*high*_), which is the maximum effective life offered by doses on the low or high sections of the ERFB contour.

These results mean we can use the Δ_*RFB*_ contour to conceptually reduce the problem of dose selection from a two-dimensional problem (choice of two doses) to a one-dimensional problem: ‘how far along the Δ_*RFB*_ contour is best?’ In other words, we have established that single resistance frequencies should be equal in the final year, and the only remaining question is: ‘how strong should the mixture be?’

On analysing the parameter scan, we find that whenever the ERFB strategy is successful, low doses are optimal (Figure 6D/Table 6). To define ‘low doses’ we split the contour into 3 based on the dose sum between the minimum and maximum *viable dose sums* (acceptable yields and dose≤ 1) along the contour. There are a small number of cases for which high doses perform equally well, but for the vast majority of points in Figure 6D lower doses are preferable (and in no case are high doses better).

**Table 6.** We split the Δ_*RFB*_ contour into 3 depending on the dose sum, splitting evenly into thirds between the minimum and maximum viable values on the contour. Minimum values are determined by the lowest doses to give acceptable yields, and maximum are determined by the largest dose sum such that both doses are less than or equal to 1. Low strength mixtures were optimal in 99% of cases, meaning their effective life was at least as good as the maximum value obtained on the 51 by 51 grid. There are 5 (i.e. 1% of cases) in which there were ERFB was not optimal (as explained in Supplementary Text 5), so low strength mixtures were optimal in every case in which ERFB was optimal. Note that in some cases there were optimal values in more than one segment of the contour, meaning that these percentages do not sum to 100. In only 3.8% of cases did high strength mixtures perform optimally (and in these cases low doses were also optimal).

| Strategy | Optimal runs | Optimal runs (%) |
| --- | --- | --- |
| Low strength | 495 | 99% |
| Medium strength | 90 | 18% |
| High strength | 19 | 3.8% |

So the optimal strategy when initial resistance frequencies vary is to pick low doses along the equal resistance in breakdown year contour. These doses are often very close to minimal, but usually fractionally stronger than weakest possible acceptable mixtures. This ensures that small losses of control due to small increases in resistance are insufficient to push the strategy below the 95% threshold, as we saw in Figure 3D.

## Discussion

Optimising deployment of fungicides in order to delay spread of fungicide resistant strains remains a major challenge (***Cunniffe et al., 2015a***). Previous work has indicated that optimal mixtures of pairs of fungicides which are both at a high risk of resistance can be constructed by using pairs of doses which select equally for both single resistant strains in the first year of application (***Hobbelen et al., 2013***). However, we have shown that this recommendation can give sub-optimal results in the common real-world case in which the initial levels of resistance to the chemicals are not equal (Figures 3, 4). We have presented an alternative strategy which gave optimal results essentially all of the time across a broad scan (Tables 4, 5, Figure 6). This was an improvement when compared to the existing strategy recommendation which was optimal 72% of the time, where we tested a range of epidemiological and fungicide efficacy parameters, as well as different initial conditions for the proportion of single- and double-resistant pathogen strains. When the initial single resistant frequencies differed by at least 10^−4^, the mean difference in effective life for the strategies was 1.01 years with a range of 0 to 4 years and the mean effective life of the equal resistance frequencies at breakdown (ERFB) strategy was 10.75 years. The average improvement in this case (as a percentage of the optimal ERFB effective life) was 7.61% with a range of 0 to 28.57%.

The strategy which consistently gave optimal results required doses which make the single resistance frequencies equal in the breakdown year. That means that the levels of resistance to each fungicide in the mixture are equal in the year that the yield becomes unacceptable due to pathogen evolution. This concept was shown to work even when fungicide parameters (asymptote, curvature and decay rate) and pathogen parameters (initial frequencies and between-season sexual reproduction proportion) were varied (Tables 4, 5). The equal resistance frequency at breakdown strategy can be framed in biological terms rather than the purely mathematical framework relied on by a naive grid-search method. This leads to the following general and practically applicable recommendations:

- reduce doses of fungicides which have higher levels of existing resistance;
- lower doses are preferable to higher;
- relatively lower doses of the higher efficacy fungicide in the mixture are preferable;
- higher efficacy could be in terms of: the decay rate (slower decaying); the curvature (the effectiveness increases more rapidly with dose); or the asymptote (the maximum level of control for that chemical).

The first point means that equal selection in the first year (ESFY) is only optimal if the initial resistance levels are sufficiently close, or equal as in ***Hobbelen et al***. (***2013***). If they differ, the mixture should be adjusted such that selection is weaker for the fungicide with an increased level of existing resistance, such that the two single-resistant strains reach equal frequency as the mixture becomes unable to achieve acceptable levels of control. Conversely, in the limit as initial resistance to one fungicide decreases, its optimal dose increases towards full dose and the dose of the mixing partner decreases to minimal levels required to give control. This matches the recommendation given by ***Elderfield et al***. (***2018***) about mixtures of low-risk and high-risk fungicides. Although the idea of low doses and relatively lower doses of higher efficacy fungicides have been presented before (***Hobbelen et al., 2013***), they have only previously been tested in the absence of pathogen sexual reproduction. We have confirmed that these recommendations also apply when there is between-season sexual reproduction (Figure 6D).

The requirement that single resistance frequencies were equal at breakdown defines a curve of possible dose pairings in dose space (the ERFB contour, see Figures 3, 4). This contour starts from a minimum viable mixture strength and runs to a maximum viable strength in dose space due to legal limitations on doses i.e. the point on the contour at which the dose of either fungicide first reaches 1. The parameter scan showed that low doses along this contour were optimal in virtually every case (Table 6, Figure 6D). The 5 exceptions are described in Supplementary Text 5 and were based on an edge effect caused by the 95% yield threshold - meaning that in these cases greater control was achieved in the final year by higher doses of either fungicide *A* or *B* despite slightly worse resistance management.

The parameter scan results can be extended to a wider range of fungicides than explicitly tested in the scan. This is because there is a mathematical link between the curvature and dose parameters which means two fungicide/dose combinations are equivalent if the curvature multiplied by chemical concentration is constant. This means that two fungicides with the same asymptote and decay rate parameters, but different curvatures can be treated as identical with an appropriate scaling of dose – see also Supplementary Text 4). This means that the ERFB strategy is optimal across many more choices of the curvature parameters than were explicitly tested in the scan.

Although sexual reproduction has a major role in the population genetics of Septoria (***Chen and McDonald, 1996***; ***Singh et al., 2021***), it has been omitted from previous models of fungicide resistance (***Hobbelen et al., 2013***; ***Elderfield et al., 2018***), justified because Septoria spreads largely clonally within season. We use the simplest model of sexual reproduction, that mating is fully random between unlinked loci and only occurs between seasons. Although loci may be linked for genes which are close together on the same chromosome, resistance of Septoria to some high-risk fungicide classes is conferred by mutations to genes on different chromosomes, leading to independent assortment. For example, resistance to methyl benzimidazoles are caused by mutations to chromosome 1 whereas resistance to succinate dehydrogenase inhibitors can be caused by mutations to chromosome 4, 7 or 8 (***Hartmann et al., 2020***). The proportion of sexual reproduction between seasons is assumed to be constant, despite evidence that sexual reproduction is influenced by density of infection (***Suffert et al., 2018***; ***Eriksen et al., 2001***). We showed that changing the between-season sexual reproduction proportion had an effect on the maximum effective life (Table 3, Figure 5). However, we showed that the inclusion of sexual reproduction did not affect the optimal strategy recommendation for a variety of different values of the proportion of between-season sexual reproduction (Table 5). Further, the model already normalises for density each year in the constant inoculum assumption which was used in previous models (***Elderfield et al., 2018***; ***Hobbelen et al., 2013***). Although the model neglects within-season sexual reproduction, we explored scenarios where all between-season reproduction was sexual, meaning that we have explored scenarios involving extensive pathogen sexual reproduction. Our results apply to haploid pathogens, although the situation may be more nuanced in the diploid case. The insecticide literature suggests that higher doses of insecticide mixtures can be preferable in tackling resistance in sexually reproducing diploid pest species, particularly in combination with refugia (***Andow and Zwahlen, 2006***).

The question of mixtures vs alternations is still under debate (***Corkley et al., 2021***), but we showed that high-risk mixtures outperform alternations when between-season reproduction occurs (Supplementary Text 3), which has previously only been shown in the case without pathogen sexual reproduction (***Hobbelen et al., 2013***).

Although these results apply to the wheat-Septoria pathosystem, they could potentially transfer to other pathosystems that are managed by mixtures of high risk fungicides. Future work may consider extending these ideas to such pathosystems e.g. grapevine powdery mildew (***Elderfield et al., 2018***) or potato blight (***Pacilly et al., 2018***; ***Cohen and Rubin, 2020***). However, we also note that past research on mixtures containing a high risk fungicide and a low risk fungicide (***Elderfield et al., 2018***) showed a remarkable consistency of optimal recommendation across pathosystems and model structures, and we have no reason to believe results would differ for other pathosystems.

The same ideas should generalise to mixtures of three or more fungicides - all modes of action in the mixture would need balancing in a similar manner. However, further work would be required to confirm this and to explore other strategies possible with more fungicides such as alternations with three chemicals or varying pairs of two chemicals. Although growers use three-way fungicide mixtures (***Phelan, 2017***), to the best of our knowledge, no modelling study exists of a mixture of three or more fungicides as applied to Septoria. Three-way fungicide mixtures were also tested for their control of *Phytophthora infestans* on tomato or potato and *Plasmopara viticola* on grape (***Samoucha, 1987***). Combining our results with those of ***Elderfield et al***. (***2018***) would suggest that a full dose of a low-risk chemical combined with minimal doses along the ERFB contour is likely to produce the longest effective life for a three-way mixture containing two high-risk fungicides and one low-risk fungicide (applied to Septoria).

In common with all theoretical studies, a number of assumptions were made in the model used to generate these results. For example, we do not consider spatial effects (***Shaw, 2000***; ***Parnell et al., 2005, 2006***), which could explore how resistance might develop on a regional scale. Management decisions made by one grower affect the entire community due to dispersal of aerial inoculum (***Laranjeira et al., 2020***). Future work might address how decisions made by growers can adversely affect others, and what policy changes can be used to incentivise sustainable behaviour. A spatially explicit model would also allow reactive disease management to be optimised (***Cunniffe et al., 2015b, 2016***), which would correspond to levels of fungicide treatment depending on local disease intensity, or even the results of a decision support type system (***Carisse, 2010***; ***Jorgensen et al., 2017***; ***Lázaro et al., 2021***).

Emergence of new resistant strains is not considered (***Hobbelen et al., 2014***), nor is partial resistance (***Mikaberidze et al., 2017***; ***Hobbelen et al., 2013***), cross-resistance (***Rehfus et al., 2018***) or fitness costs (***Mikaberidze et al., 2014***; ***Mikaberidze and McDonald, 2015***). Strong fitness costs to resistance could prolong effective lives by reducing the overall benefit of carrying resistance. If the double-resistant strain carried the strongest penalty, we would expect the ERFB strategy to still hold since the single resistant strains become a relatively greater threat to control (and the ERFB strategy performs well at managing the single resistant strains). Similarly inclusion of partial resistance would further extend effective lives, since then resistant strains would be suppressed to some extent by the fungicide. We believe that the ERFB strategy would still hold in this case; future work would be needed to confirm this.

There is no stochasticity in the model, so variation within a growing season and between different growing seasons is ignored (***te Beest et al., 2008***), as is precise timing of fungicide applications (***van den Berg et al., 2013, 2016***). Effects of host plant resistance (***Mikaberidze and McDonald, 2020***; ***Carolan et al., 2017***) and of multiple pathogens (where the crop is infected by e.g. brown rust as well as Septoria (***Garin et al., 2018***)) are neglected. These various factors potentially have effects on the strength of mixture required, although different relative dosages of the two fungicides could still be used to give greater protection to the chemical at most urgent threat.

Future work might focus on exploring total lifetime yield (***Elderfield et al., 2018***) or economic output (***te Beest et al., 2013***; ***van den Bosch et al., 2020***) rather than effective life. The latter would involve incorporating the price of grain into the calculations, as well as the cost of each fungicide application and other costs from each year’s farming. Using economics rather than effective life would likely incentivise lower doses in general, and relatively higher doses of cheaper fungicides if the difference in price relative to efficacy was significant. For instance, if a cheap high efficacy fungicide were used, lower doses might be best for effective life, but higher doses might lead to cost savings within an individual year. In a system where grain prices are high, each additional year of relatively high yield is very valuable, so resistance management becomes the most important factor. This means economics would probably only change where it is optimal within the small region of dose space which contains the maximum effective life. In a system where grain prices were relatively low, then minimising doses of expensive fungicides may become a higher priority than choosing a strategy which is optimal for resistance management. These factors are important when considering policy changes to incentivise sustainable use of fungicide mixtures. Use of modelling to inform policy change on sustainable agriculture should be informed by agronomic, environmental, economic, and social considerations (***Mouratiadou et al., 2021***).

The breakdown in the final year recommendation may seem less practical than a first year recommendation, since it may be difficult to predict the future evolution of the pathogen population, leading to greater uncertainty in the recommendation. However, points in dose space that are close to the optimal ones usually also have a long effective life (Figure 4). This means that even if the estimates of the initial resistance frequencies or other parameters are imperfect, a good decision that is close to the optimal can be made. The model could be used to find the best estimate for the optimal dose-pairing given imperfect information about the levels of resistance or fungicide parameters. It can be difficult to estimate the proportion of resistance strains particularly when their incidence is very low (***van den Bosch et al., 2014a***) and there is regional variation (***Garnault et al., 2019***). However, as the resistance frequencies increase and become easier to reliably estimate, the model output could be updated with an improved estimate for the corresponding optimal dose combination. As is invariably the case, more complex disease management recommendations (such as the equal resistance at breakdown recommendation) require good prior knowledge for good outcomes (***Hyatt-Twynam et al., 2017***).

There are some practical criticisms which can be made – for example, growers typically apply doses in multiples of a quarter of a full dose. This means that a very precise theoretical prescription for an optimal dose may not be used under current practices. However, our results show that pairs of fungicides should be used in such a way as to avoid resistance developing too rapidly to either component of the mixture. This may be achievable to an extent even if we restrict our attention to multiples of quarter-doses. Further, if modelling shows that using more precise doses would result in dramatic increases in the durability of a particular fungicide mixture then they should be recommended.

An interesting area for future work would be to consider time varying disease management strategies. Given that the resistant frequencies vary each year, it may be possible to prolong the effective lifetime by increasing dose as the level of resistance increases (and the reliability of estimates resistance frequency improves). Further, it would allow other strategies like alternating the use of mixtures that favour fungicide *A* or *B*, which could outperform strategies that are static in time. We hope to address these questions in future work. One approach to this would be to use optimal control theory (***Bussell et al., 2019***; ***Bussell and Cunniffe, 2020, 2022***), another would be to use dynamic programming (***Bellman, 1952***; ***Onstad and Rabbinge, 1985***).

## Availability of code online

An implementation of the model in the freely-available programming language Python (Python Software Foundation, available at http://www.python.org) is online at https://github.com/nt409/HRHR.

## Acknowledgements

NPT acknowledges the Biotechnology and Biological Sciences Research Council of the United Kingdom (BBSRC) for support via a University of Cambridge DTP PhD studentship (Project Reference 2119688). The authors also thank Alexey Mikaberidze, Christopher Gilligan and Julia Davies for useful discussions.

## Supplementary Text 1

### Between-season sexual reproduction – derivation

For between-season sexual reproduction, we assume that the recombination is random, so that each (haploid) offspring has an equal probability of inheriting each allele from either (haploid) parent (Supplementary Text 1 Table 1). We have two unlinked (assumed) loci, each with two alleles: resistant or sensitive.

**Supplementary Text 1 Table 1.** The possible offspring of sexual reproduction between parents given by the left column and top row are shown. For those entries which contain four strains, the probability of each is 1/4, so each number is accordingly scaled. Similarly those entries containing two strains each have probability 1/2. Note that both parents and offspring are haploid; for example *sr* corresponds to a haploid which carries the allele for sensitivity (*s*) to fungicide *A* on one chromosome, and the allele for resistance (*r*) to fungicide *B* on another.

| Pathogen strains | *ss* | *sr* | *rs* | *rr* |
| --- | --- | --- | --- | --- |
| *ss* | *ss* | *ss/sr* | *ss/rs* | *ss/sr/rs/rr* |
| *sr* | *ss/sr* | *sr* | *ss/sr/rs/rr* | *rr/sr* |
| *rs* | *ss/rs* | *ss/sr/rs/rr* | *rs* | *rr/rs* |
| *rr* | *ss/sr/rs/rr* | *rr/sr* | *rr/rs* | *rr* |

Let *F*_*mn*_ denote the frequency (a proportion ∈ [0, 1] of total disease) of strain *mn* immediately before the single, instantaneous round of between-season sexual reproduction which we model. Let *Y*_*mn*_ represent the frequency of strain *mn* immediately afterwards. Then collecting together all the terms, we find:

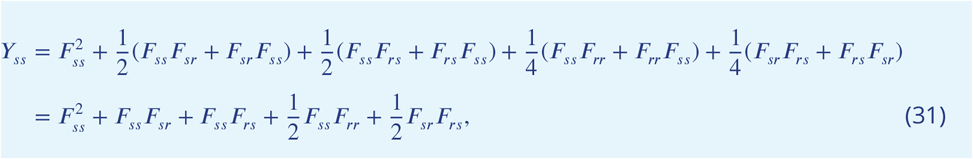

and similarly:

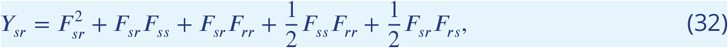

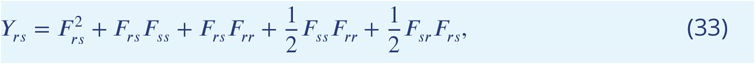

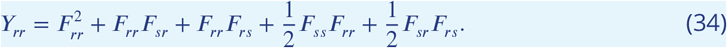

This can be simplified by noting that, *F*_*ss*_ + *F*_*sr*_ + *F*_*rs*_ + *F*_*rr*_ = 1, and that, using the *ss* strain as an for example:

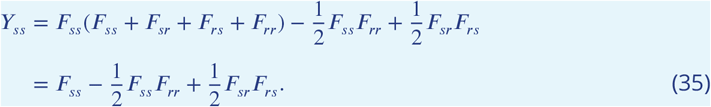

Now define:

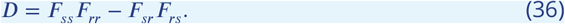

Then we find:

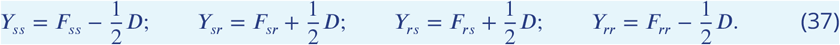

This form is equivalent to that of ***Felsenstein*** (***1965***) for a haploid population, with two alleles at two loci and discrete generations (we use *F*_*mn*_ in place of their *x*_*i*_). We assume that the recombination fraction *r* = 1/2, corresponding to the two loci being unlinked. We also assume that the probability of surviving from meiosis to fertilisation is equal for all strains, in line with our assumption that the strains behave identically in the absence of fungicide.

## Supplementary Text 2

### Between-season sexual reproduction – effect on the model

Depending on initial resistance frequencies and fungicide parameterisations, we can observe:

- non-monotonic response in maximum effective life as *q*_*B*_ increases (Supplementary Text 2 Figure 1A);
- no effect on maximum effective life as *q*_*B*_ increases (Supplementary Text 2 Figure 1B);
- monotone increase in maximum effective life as *q*_*B*_ increases (Supplementary Text 2 Figure 1C).

**Supplementary Text 2 Figure 1.**
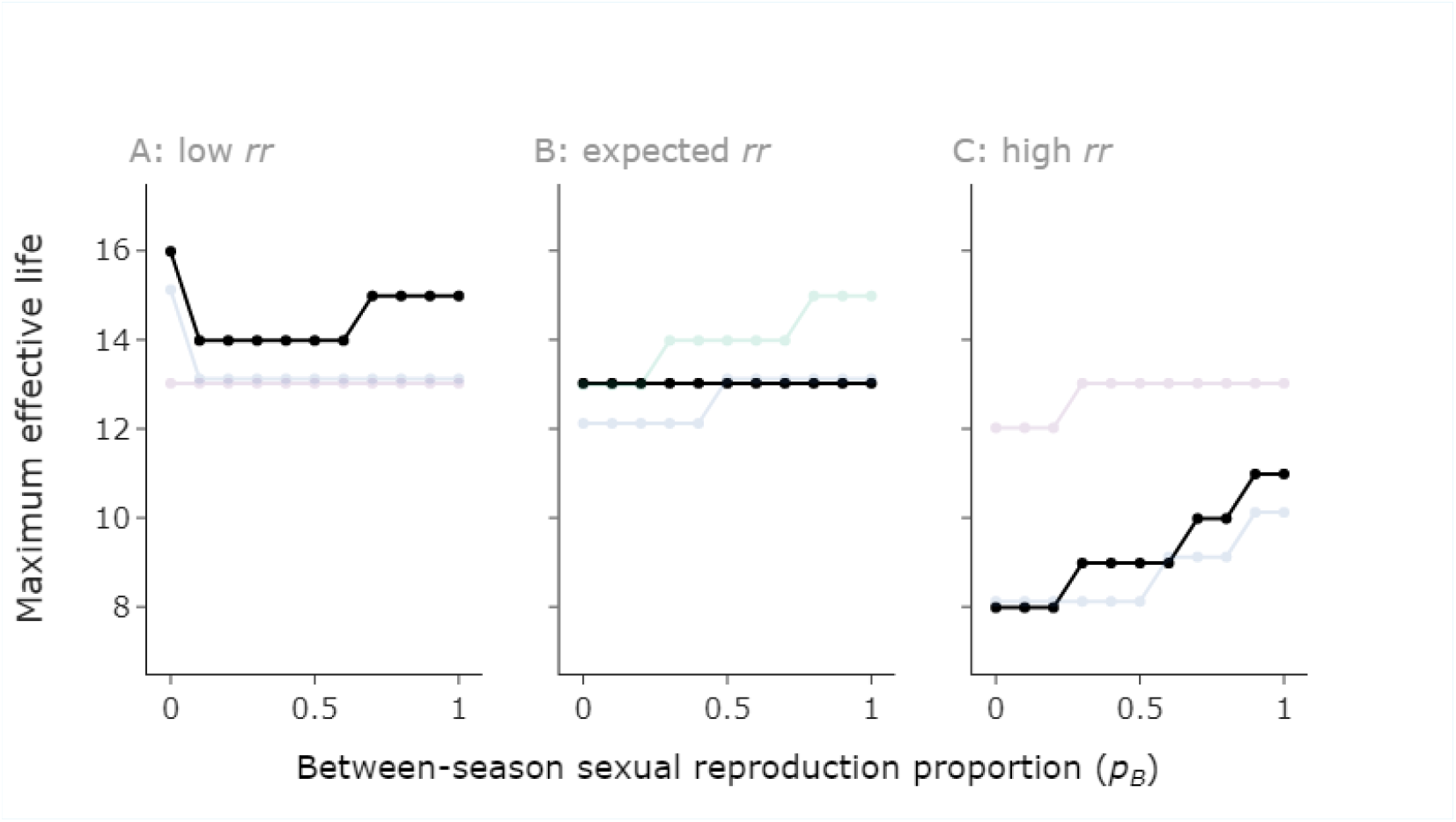
Three effects are possible with sexual reproduction. This figure is exactly as in Figure 5 (main text), but highlighting the three qualitative behaviours we see as the proportion of sexual reproduction (*q*_*B*_) increases. These are: (**A**) non-monotonicity; (**B**) no change in effective life; and (**C**) monotonic increase in effective life as *q*_*B*_ increases. **Parameter values:** as in Figure 5 (main text); random fungicide parameter values and single resistant frequencies, generated from distributions described in Table 3. **A**: initial double resistant strain (*rr*) density 10^−4^ times lower than value from linkage equilibrium (VLE); **B**: initial *rr* density at VLE; **C**: initial *rr* density 10^4^ times higher than VLE. Values: (*p*_*rr*_, *p*_*rs*_, *p*_*sr*_, Λ_*A*_, Λ_*B*_, *ω*_*A*_, *ω*_*B*_, *θ*_*A*_, *θ*_*B*_) = **Scenario 1**: (7.049916 × [10^−19^/10^−15^/10^−11^], 5.545997 × 10^−7^, 1.269711 × 10^−8^, 0.009758, 0.022030, 0.729797, 0.661193, 7.362942, 6.642679); **Scenario 2**: (1.085892 × [10^−15^/10^−11^/10^−7^], 1.596583 × 10^−6^, 6.801290 × 10^−6^, 0.007417, 0.009834, 0.574543, 0.706497, 11.143576, 11.170345); **Scenario 3**: (2.445955 × [10^−15^/10^−11^/10^−7^], 7.155682 × 10^−5^, 3.417953 × 10^−7^, 0.005803, 0.006279, 0.875035, 0.717337, 8.544356, 11.404773).

For medium or high initial levels of the double resistant, we typically only see the latter two behaviours (no change or monotone increase). This is because sexual reproduction acts to suppress the double resistant strain, as shown by the between-season equation for the *rr* strain:

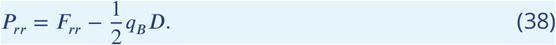

Therefore if *D* = *F*_*rr*_*F*_*ss*_ − *F*_*sr*_*F*_*rs*_ > 0, we expect a decrease in the level of double resistant after the sexual reproduction step (for any *q*_*B*_ > 0, i.e. any non-zero amount of sexual reproduction). Note that *D* > 0 is equivalent to *F*_*rr*_ > *F*_*sr*_*F*_*rs*_/*F*_*ss*_. Initially *F*_*ss*_ ≈ 1, so the threshold *F*_*sr*_*F*_*rs*_/*F*_*ss*_ ≈ *F*_*sr*_*F*_*rs*_. The effect of sexual reproduction decreasing the rate of the double resistant strain increasing is particularly clear in Supplementary Text 2 Figure 2H, and is present but to a lesser extent in Supplementary Text 2 Figure 2D.

When the effective life is the same across different levels of sexual reproduction, this is caused by increases in the single-resistant strains relative to the asexual case (Supplementary Text 2 Figure 2E,F,I,J). This offsets the decrease in the double-resistant strain. Whether the single or double resistant strain plays the major role in yield losses depends on the efficacy of the fungicides – very high efficacy fungicides are capable of controlling the single resistant strains since one component of the mixture remains effective. For example, relatively good control of the single resistant strains in Supplementary Text 2 Figure 2A,B,C means that the effective life is quite long, since it takes a long time for the double resistant to reach sufficiently high frequencies to reduce yields below 95%. In contrast, poor control from fungicide A and good control from fungicide B in Supplementary Text 2 Figure 2D,E,F means that the strains resistant to fungicide B (strains *sr, rr*) rapidly increase in density and lead to loss of control whilst strain *rs* declines.

If the level of double resistant is very low (*F*_*rr*_ < *F*_*sr*_*F*_*rs*_/*F*_*ss*_), then *D* < 0 and the double resistant increases in density (Supplementary Text 2 Figure 2A). This increase is only seen in the first year, after which point the double resistant remains at a level at least as high as *F*_*sr*_*F*_*rs*_/*F*_*ss*_, due to the greater selection for the *rr* strain in comparison to the other strains every year. The non-monotonic effect is due to even low levels of sexual reproduction leading to an increase in the double resistant after the first off-season, but higher levels of sexual reproduction again suppressing the double resistant strain in subsequent years. This is shown by the decreased gradient of sexual proportion *q*_*B*_ = 1 vs sexual proportion *q*_*B*_ = 0.5 in Supplementary Text 2 Figure 2A.

**Supplementary Text 2 Figure 2.**
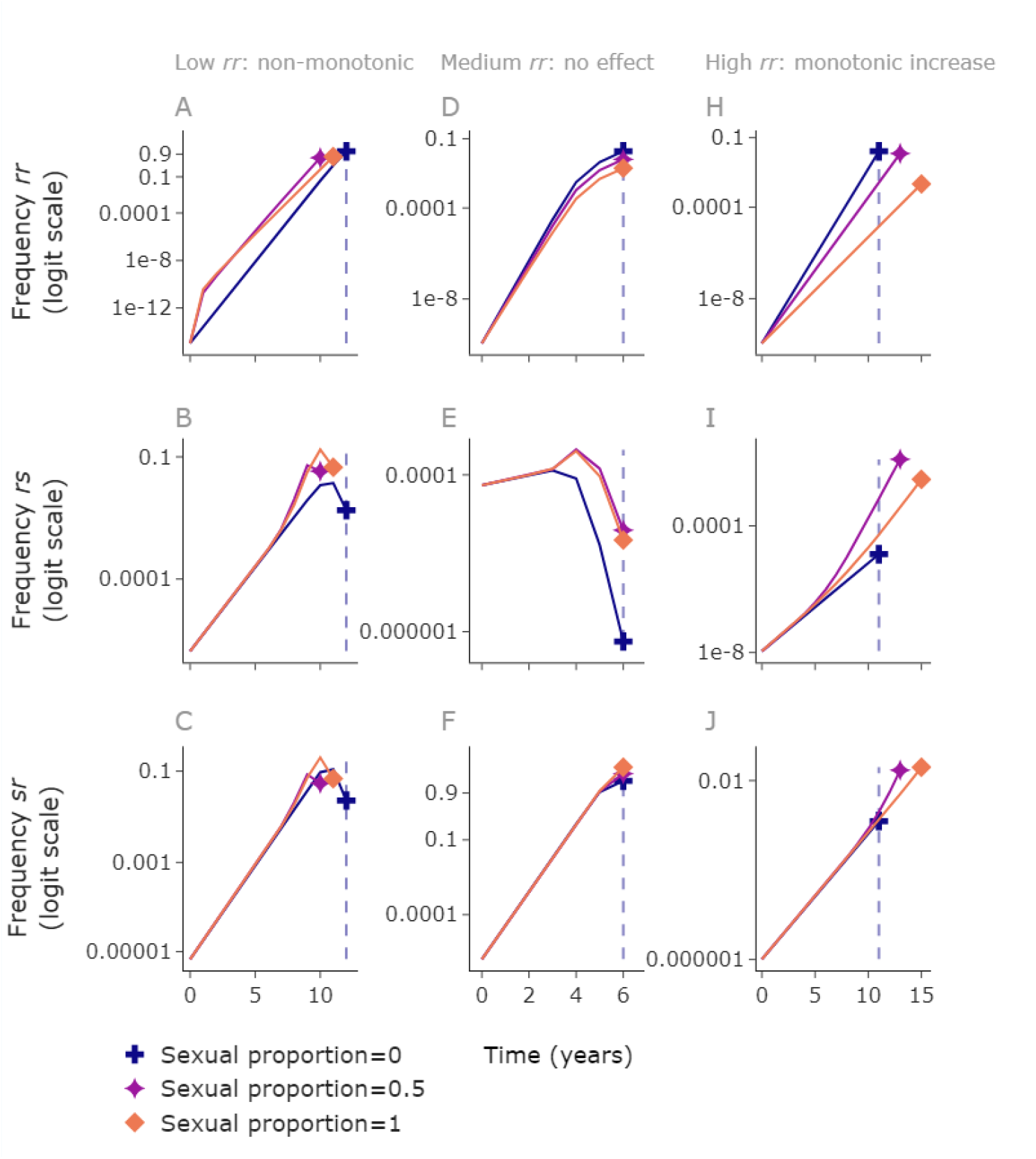
Higher between-season sexual reproduction slows the double resistant strain. Three different scenarios, demonstrating the three qualititative behaviours from Supplementary Text 2 Figure 1: non-monotonicity in effective life as *q*_*B*_ increases; no change in effective life as *q*_*B*_ increases; monotonic increase as *q*_*B*_ increases. The dotted line is the effective life of the asexual reproduction case. Note that here we compare full dose of both fungicides in each case, whereas in Supplementary Text 2 Figure 1 and main text, Figure 5, we compare the optimal dose from the 21 by 21 grid of doses in each case. The randomly generated scenarios that we show here are different to any shown in Supplementary Text 2 Figure 1, but here we are comparing something different (full dose rather than optimal dose). **NB** the subplots in each column are for the same scenario, but the scenarios are completely different between columns (not just a change in initial resistance frequencies). **Left column**: the ‘non-monotonicity’ case. In this example the double resistant strain is lower than the value at linkage equilibrium (VLE) by 10^−4^. Here high values of *q*_*B*_, or *q*_*B*_ = 0 give longer effective lives than intermediate values. **Middle column**: the ‘no effect’ case. In this example the double resistant strain is at VLE. Here there is a faster increase in the double resistant strain when *q*_*B*_ = 0 (**D**) but this is offset by reduced growth of the single resistant strains (**E**,**F**). **Right column**: the ‘monotone increase’ case. In this example the double resistant strain is higher than the VLE by 10^4^. Here the faster growth of the double resistant strain when *q*_*B*_ = 0 causes a shorter effective life than we see in the case where *q*_*B*_ takes larger values. **Parameter values**: Generated at random: (*I*_*rr*_, *I*_*sr*_, *I*_*rs*_, *ω*_*A*_, *ω*_*B*_, *θ*_*A*_, *θ*_*B*_, Λ_*A*_, Λ_*B*_) = **left column**: (1.085892 × 10^−15^, 6.8012899586885546 × 10^−6^, 1.596583402894029 × 10^−6^, 0.574543, 0.706497, 11.143576, 11.170345, 0.007417, 0.009834); **middle column**: (1.140463 × 10^−10^, 1.545001824789278 × 10^−6^, 7.38107054353044 × 10^−5^, 0.983611, 0.82889, 9.581831, 5.728716, 0.032598, 0.003884); **right column**: (1.121325 × 10^−10^, 1.0174155443038875 × 10^−6^, 1.1002632709098869 × 10^−8^, 0.697464, 0.480298, 5.136889, 5.748469, 0.016088, 0.011044).

## Supplementary Text 3

### Between-season sexual reproduction – mixtures vs alternations

We now explore the effect of varying the between-season sexual reproduction *q*_*B*_ on the model output and the optimal strategy. To check that mixtures outperform alternations when between-season sexual reproduction was present, we ran a simple scan where between-season reproduction was entirely sexual. The case where reproduction was entirely asexual has previously been examined in ***Hobbelen et al***. (***2013***); ***Elderfield et al***. (***2018***), and mixtures were found to outperform alternations.

We tested 100 randomly generated scenarios (Supplementary Text 3 Table 1). We varied the asymptote, curvature and decay rate parameters of both fungicides, as well as the initial resistance frequencies, ignoring any parameter sets where either of the alternation strategies had an effective life of 0 (i.e. could never achieve a yield ≥ 95%). Using a grid of 51 × 51 doses, we checked whether the maximum effective life from the mixture tactic in each case was greater than or equal to the maximum effective life from either alternation tactic (applying fungicide *A* first each year or applying fungicide *B* first each year). In every case, the mixture tactic was at least as good as either alternation tactic (Supplementary Text 3 Figure 1).

**Supplementary Text 3 Table 1.**
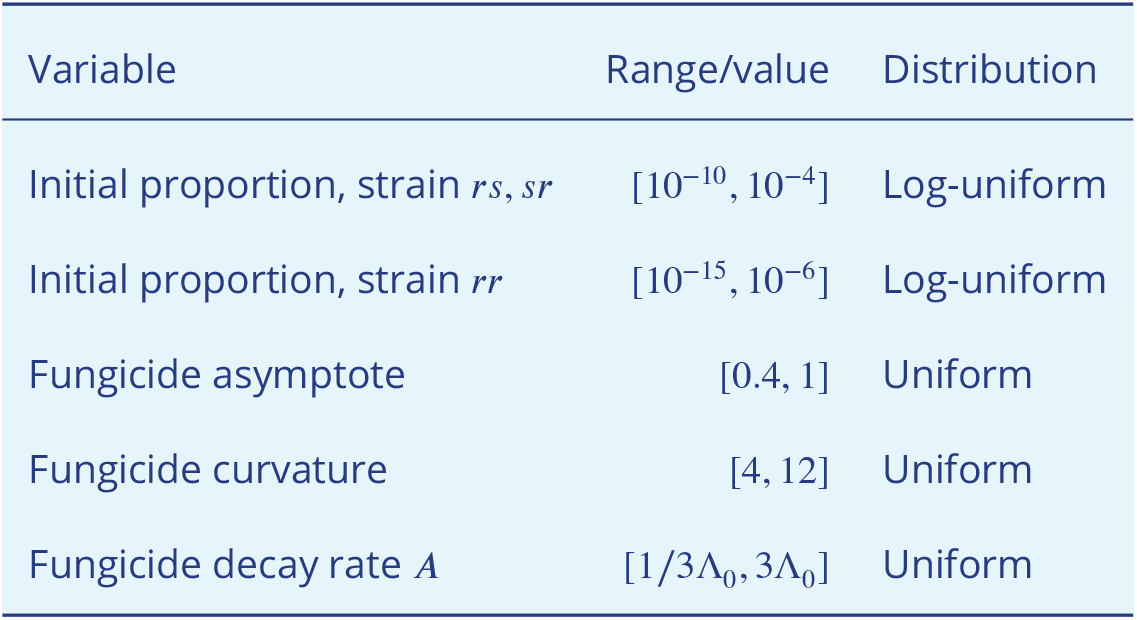
Bounds for the parameter scan used to confirm that mixtures outperform alternations. For this scan we used a grid of 51 × 51 doses, and checked that the maximum effective life from the mixture grid was greater than or equal to the maximum from the alternation grid for each example. We tested the case where between-season reproduction was entirely sexual. Unless stated above, all other parameters take their default values (see Table 1). Note the fungicide decay rate range depends on Λ_0_, which is 1.11 × 10^−2^ degree days^-1^ as in Table 1.

**Supplementary Text 3 Figure 1.**
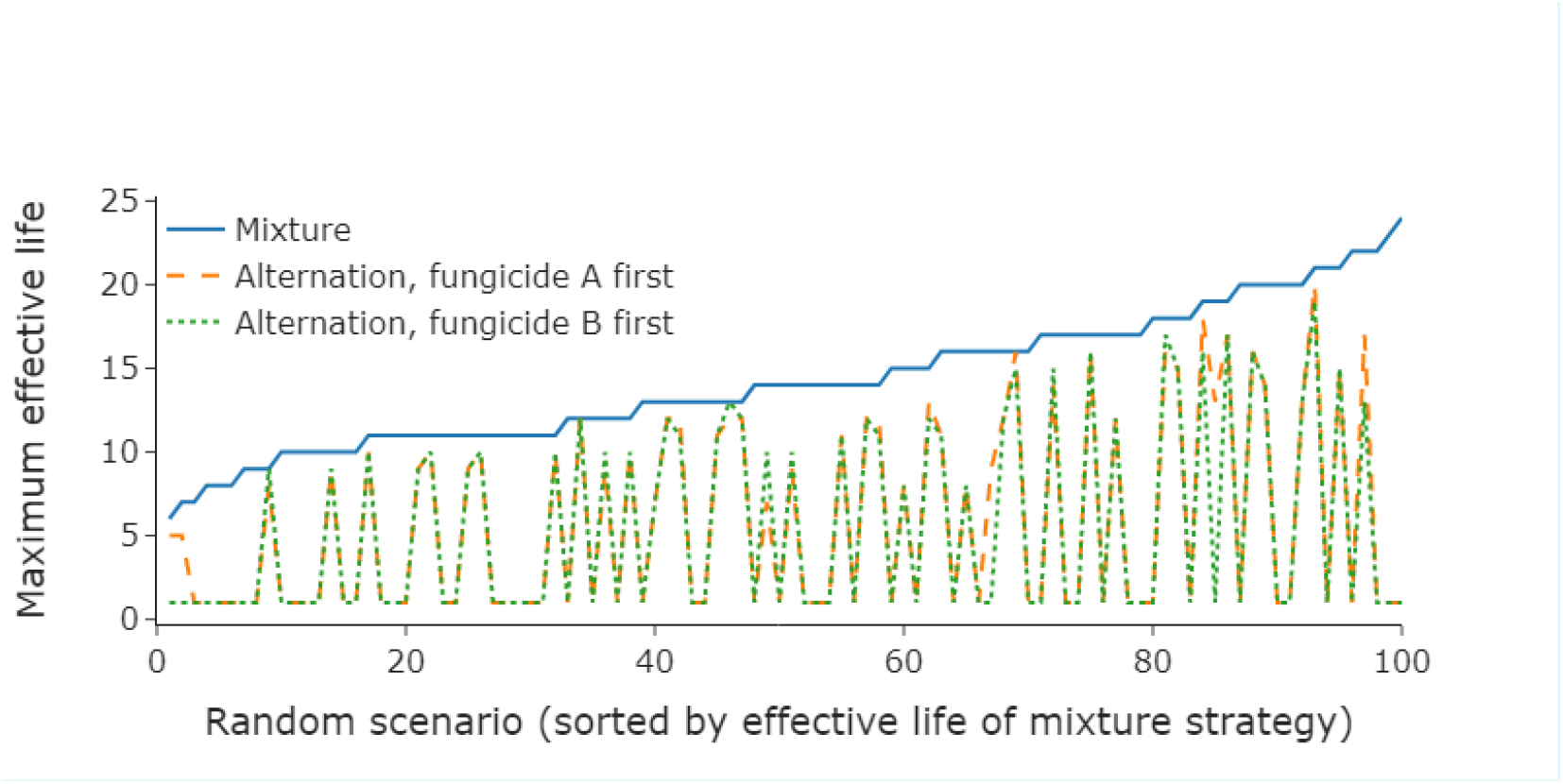
Mixtures outperform alternation even with between-season sexual reproduction. Here are the optimal effective lives from 100 randomly generated scenarios. The scenarios are arranged from smallest to largest effective life of the mixture strategy. Here we consider the optimal dose from a 51 by 51 grid. **Parameter values**: randomly generated, see Supplementary Text 3 Table 1.

## Supplementary Text 4

### Parameter scan – link between curvature and dose

The following argument will show the link between curvature and dose which allows us to generalise the parameter scan results (Tables 5, 6 and Figure 6) to a wider range of fungicides. In particular, we show that any pair of fungicides with identical decay rate and asymptote but differing curvature can be considered to behave identically with an appropriate change of dose. To meet legal dose requirements (doses less than 1), we note that this applies to any higher efficacy fungicides (increased curvature) than those tested, and some lower efficacy ones (see below).

For a dose *C* of a fungicide *F*, we use dose responses of the type

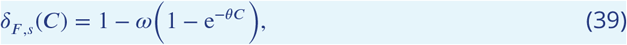

where *δ*_*F,s*_(*C*) ≤ 1 is the factor by which the rates of the transition of tissue from healthy to latent (*β*), and from latent to infected (*γ*) are reduced. Now define *ε* as the effect on sensitive strains:

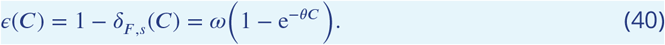

We also assume fungicide concentrations decay exponentially, so that *t* units after a dose application the concentration will be:

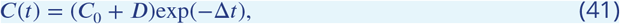

where Δ is the decay rate, *C*_0_ was the concentration immediately before the application, and *D* was the applied dose.

Consider a pair of fungicides 1 and 2 which differ only in their curvature, i.e. have parameters (*ω, θ*_1_, Δ) and (*ω, θ*_2_, Δ). Then if we apply a dose of fungicide 2 such that

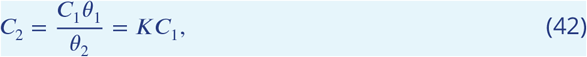

then we find:

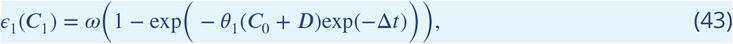

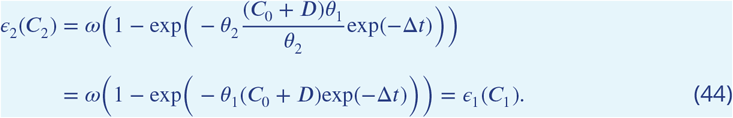

That is, if fungicide 1 has a curvature that is a factor of *K* bigger than fungicide 2, we can apply doses that are a factor of *K* smaller than fungicide 2. Then we end up with an identical form for *∈*_1_ and *∈*_2_, and so an identical form for *δ*_1,*s*_(*C*_1_) and *δ*_2,*s*_(*C*_2_), even when accounting for the decay in concentration over time. This means that the two fungicides behave identically if these doses are applied (Supplementary Text 4 Figure 1).

**Supplementary Text 4 Figure 1.**
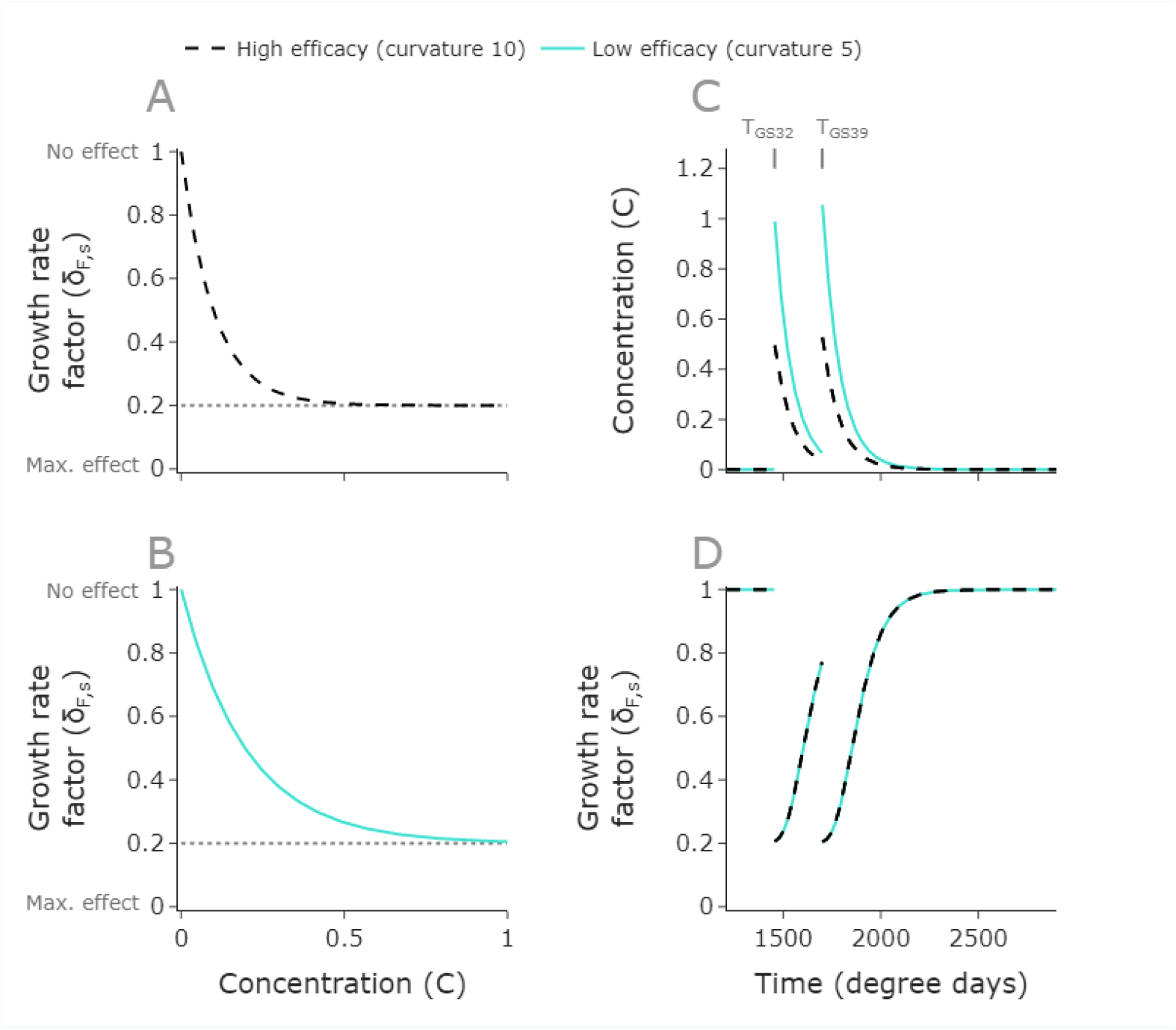
Fungicides with different curvatures can behave identically with appropriate dose choice. One fungicide has a curvature *θ*_*h*_ = 10 and the other has a curvature of *θ*_*l*_ = 5. Both have the same asymptote and decay rate, which is required for this effect. This parameterisation results in the former fungicide having a higher efficacy (**A, B**). Applying appropriately scaled doses: *D*_*h*_ = *D*_*l*_*θ*_*l*_/*θ*_*h*_, so that *D*_*l*_ = 1, *D*_*h*_ = 0.5, (**C**), the effect is identical (**D**). Since the lower efficacy fungicide has half the curvature, it requires double the dose. **Parameter values**: higher efficacy fungicide *θ*_*h*_ = 10. Lower efficacy fungicide: *θ*_*l*_ = 5. Both have asymptote *ω* = 0.8 and decay rate Λ = 1.11 × 10^−2^. Doses applied in **B**: (*D*_*h*_, *D*_*l*_ = 0.5, 1).

### Legal dose caveat

Note that to ensure that we remain within legal requirements for doses (*C* < 1), we require that

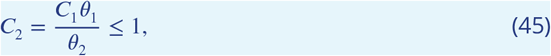

so the parameter scan analysis can only directly be transferred to fungicides with curvatures satisfying

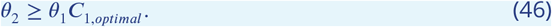

Since minimal doses tend to be optimal, this means our results may be generalised for a range of fungicides that are lower or equal efficacy than those in the scan (*θ*_1_*C*_1,*optimal*_ ≤ *θ*_2_ ≤ *θ*_1_), and any fungicides that are higher efficacy (higher curvature than *θ*_1_).

For instance, if a particular fungicide with curvature 8 had an optimal dose of 0.25 in a mixture, then we could apply our results to any fungicide with curvature *θ* ≥ 8 × 0.25 = 2, using optimal dose 2/*θ* ≤ 1.

## Supplementary Text 5

### Parameter scan – sub-optimal runs

#### Runs in which equal resistance frequencies at breakdown was sub-optimal

There were initially 8 runs of the 500 for which equal resistance frequencies at breakdown (ERFB) appeared to not give optimal results. However, when re-running so that a much larger number of doses along the contour were chosen (500 and in a narrower range around the optimum), 3 of these cases were found to be optimal, leaving only 5 runs that were sub-optimal after the more computationally intensive search. The remaining 5 runs (Supplementary Text 5 Figure 1) the results remained sub-optimal by one year even when the denser grid was used. Parameter values for these runs are in Supplementary Text 5 Table 1.

In each of these cases, the optimal region is very small. There is a much larger region which achieves an effective life within one year of the optimal value. In this larger region, the resistance management offered is relatively good. The doses in the optimal region offer better control in that year than the surrounding region. The location of this region depends on the efficacy of the two fungicides, and in these 5 cases the ERFB contour does not pass through the optimal region. We examine one of these runs (scenario B) in Supplementary Text 5 Figure 2.

Some dose-pairs near the optimal region are sub-optimal effective lives because they offer slightly poorer control, despite good resistance management. Other higher dose choices near the optimal region are sub-optimal because the poorer quality resistance management is bad enough to not offset the greater control caused by higher doses in the final year.

**Supplementary Text 5 Figure 1.**
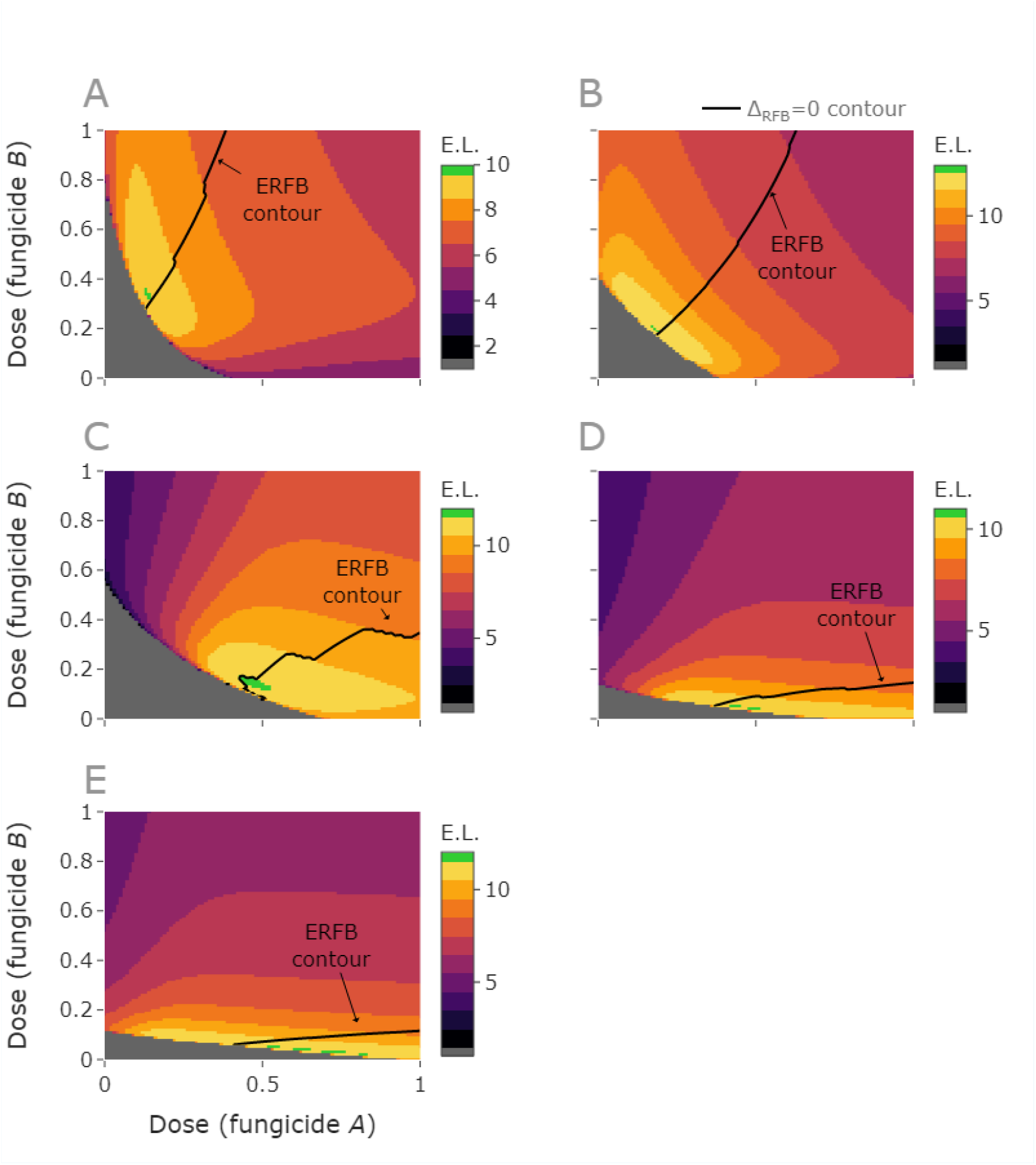
All sub-optimal runs. All 5 sub-optimal runs are shown, here on a 101 by 101 grid. In each case the optimal region is very small and gives a one year improvement relative to the best dose along the ERFB contour. In each case the contour narrowly misses the optimal region. **Parameter values**: see Supplementary Text 5 Table 1.

**Supplementary Text 5 Table 1.**
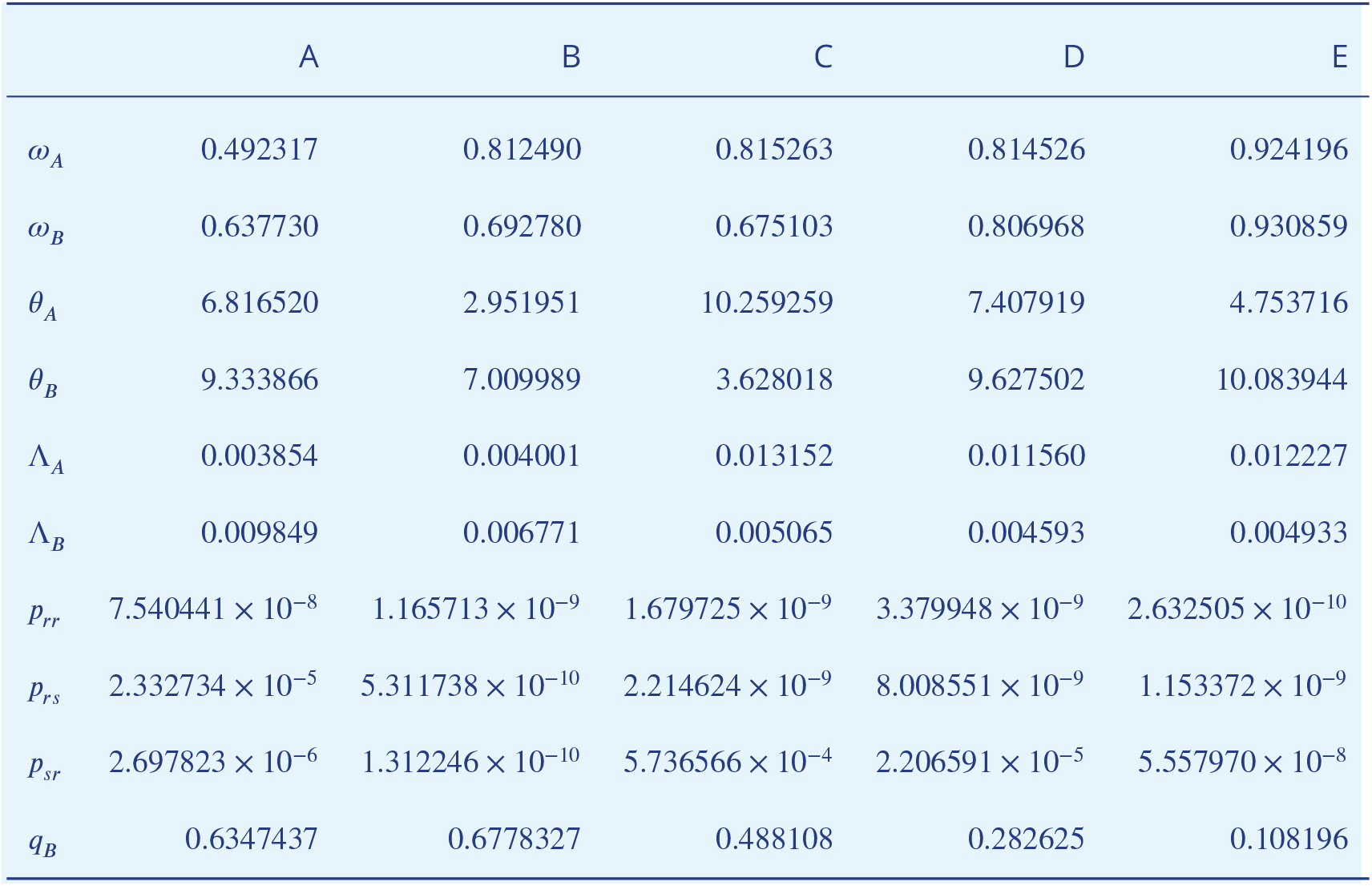
Parameter values for the sub-optimal runs shown in Supplementary Text 5 Figure 1.

**Supplementary Text 5 Figure 2.**
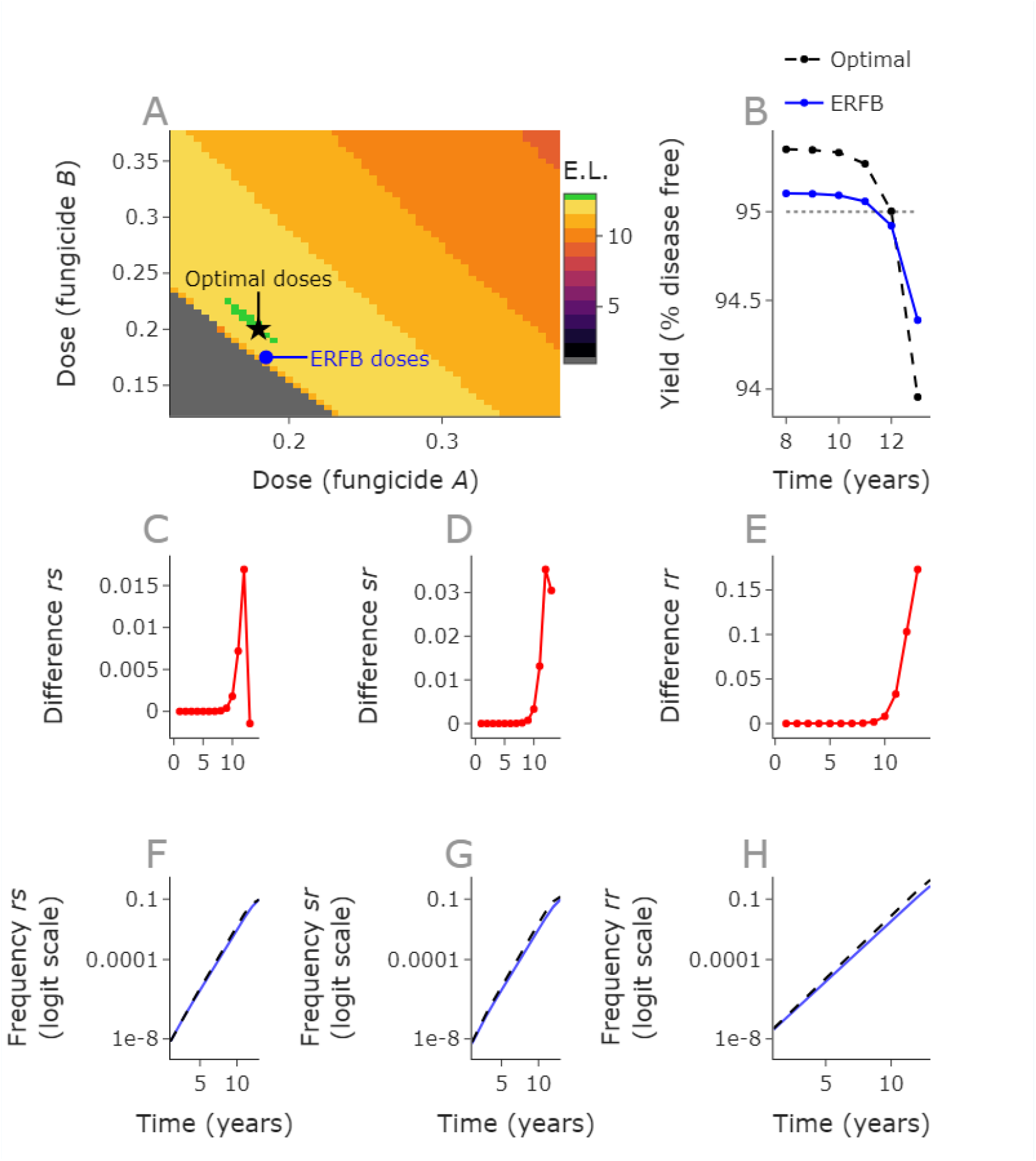
Final year control causes the small number of exceptions where the ERFB strategy is sub-optimal. The optimal strategy achieves a yield of 95.003% in year 12, compared to the ERFB strategy which achieves 94.92% in this year (**B**). The optimal strategy fails in year 13, whereas the ERFB strategy fails in year 12 (**A**). Despite lower incidences of all resistant strains under the ERFB strategy, (**F, G, H**), the control offered by the doses is lower in year 12, since the dose of fungicide *B* is much lower. Each resistant strain increases faster under the optimal strategy, meaning that the differences in frequency increase over time (**C, D, E**). **Parameter values:** see Supplementary Text 5 Table 1, scenario B. Doses: optimal = 0.18, 0.2, ERFB = 0.185, 0.175.

Taking the first of the sub-optimal examples (Supplementary Text 5 Figure 1A), we see that for a small perturbation in initial resistance frequencies this effect disappears and ERFB is optimal (Supplementary Text 5 Figure 3). This supports the conclusion that the effect is caused by control in the final year, which is why every sub-optimal run of the 500 was only worse than the optimal by a single year. By the failure year, usually resistance frequencies are increasing rapidly and yield is falling steeply (main text, Figure 2F, 2G). This is why it is rare to have such a small optimal region; more commonly all doses within a larger optimal region are further above 95% in the penultimate year and further below 95% in the breakdown year.

**Supplementary Text 5 Figure 3.**
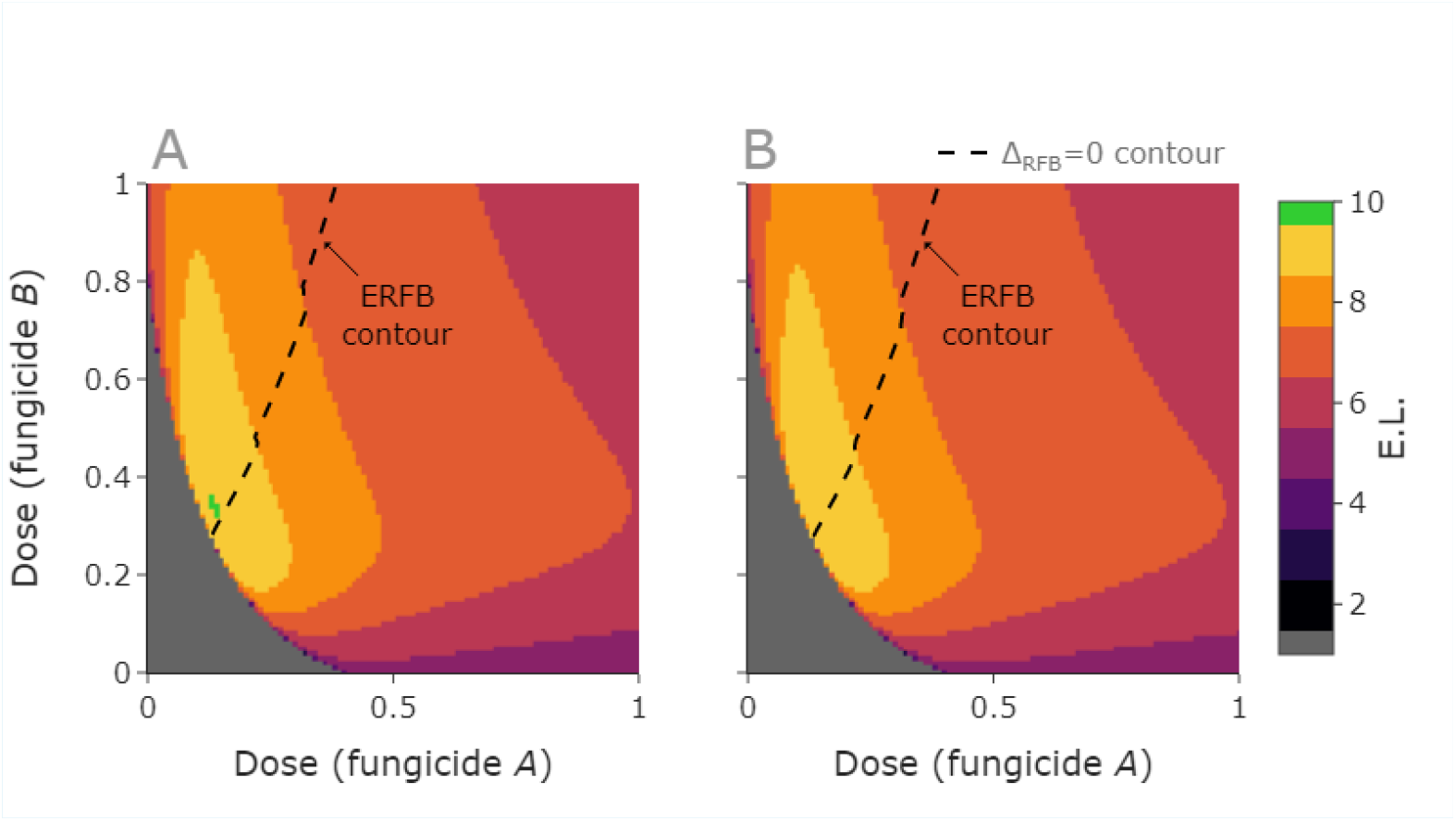
ERFB is optimal under small perturbation of initial conditions of sub-optimal run. This is the sub-optimal run with a grid size of 201 by 201. The optimal region is very small and outperforms ERFB because the increased control in offered by a higher dose of fungicide *B* (**A**). However, a small perturbation away from the initial conditions removes this effect, and we find that the ERFB strategy is optimal again (**B**). Here the green region no longer exists and the optimum effective life is now 9 (yellow). If we gradually decrease resistance frequencies, the scenario can change from something analogous to panel **B** to something analogous to panel **A**. This is because the doses which achieve yields fractionally above 95% in the final year may not be the same as those which are best for longer term resistance management, since control and resistance management may not align in any given year. In practice there are very few cases where this phenomenon is observable, as shown by the fact it does not impact 99% of cases. **Parameter values:** **Pathogen parameters, A**: see Supplementary Text 5 Table 1, scenario A. **Pathogen parameters, B**: 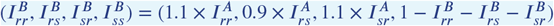.

### Run in which equal selection in the first year outperformed equal resistance frequencies at breakdown

There was a single case out of the 500 in which equal selection in the first year (ESFY) gave an improved output relative to ERFB. Here the best ERFB effective life matched the optimal value from the 51 by 51 grid, but the best ESFY effective life was one year greater than this optimal grid value (Supplementary Text 5 Figure 4). Note that Supplementary Text 5 Figure 4 uses a denser grid of 201 by 201 to adequately illustrate this effect. See also Supplementary Text 5 Figure 5. Note that the comparison in the parameter scan was between the 51 by 51 grid, ESFY and ERFB; it is possible that there are other cases where a finer grid might find small regions which achieve one year greater than the ERFB optimum and the 51 by 51 grid optimum.

**Supplementary Text 5 Figure 4.**
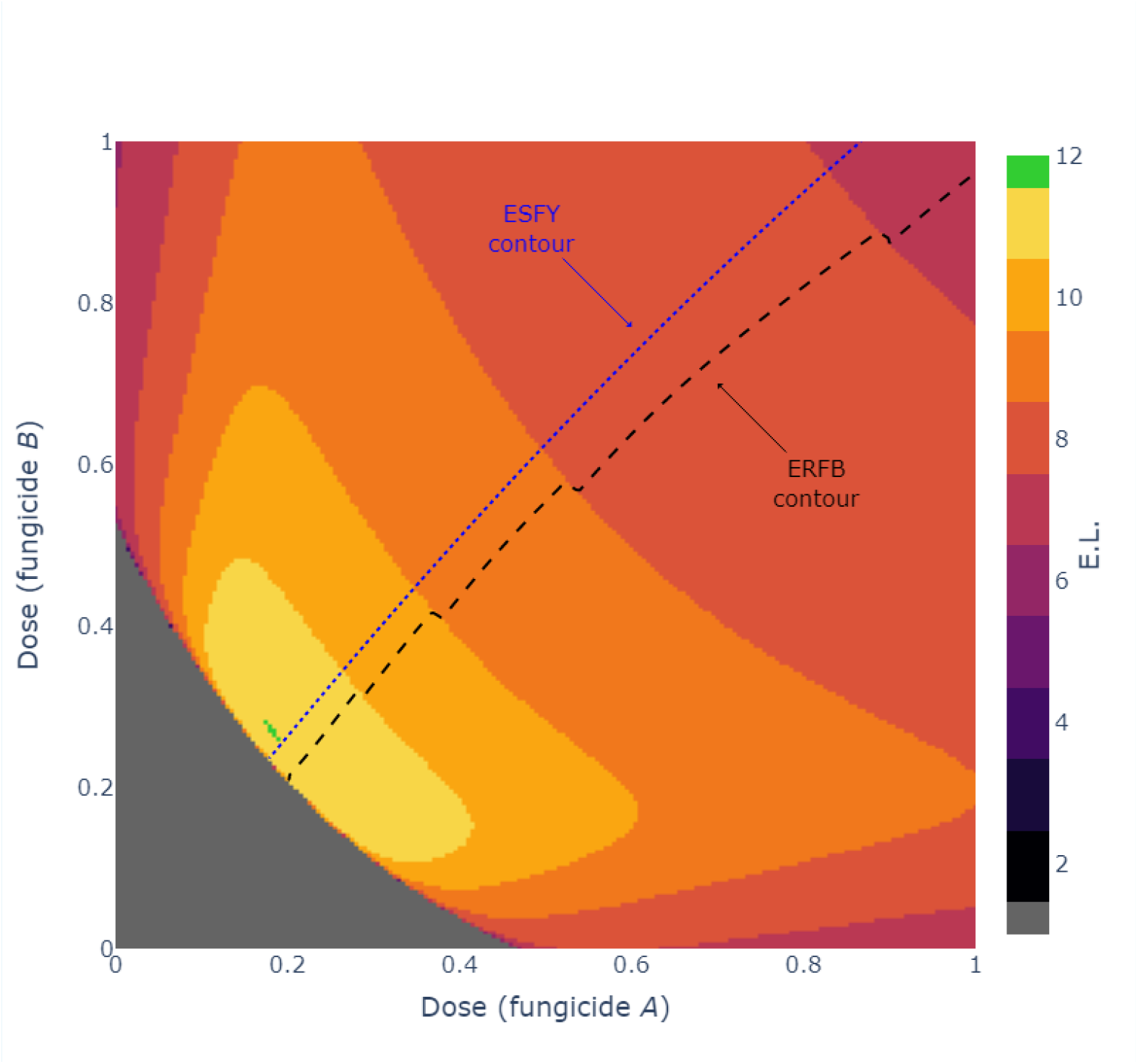
The single case where ESFY outperformed ERFB. In this case the ESFY contour passes through the optimal region, but the ERFB contour does not. This was the only case in which this happened. Although ERFB is sub-optimal on the 201 by 201 grid, it was optimal on the coarser 51 by 51 grid. **Parameter values:** Initial frequencies: *I*_*rr*_ = 1.3122930696727086 × 10^−10^, *I*_*rs*_ = 1.3687346286628245 × 10^−7^, *I*_*sr*_ = 9.216054320447214 × 10^−7^, *I*_*ss*_ = 0.9999989413898756. *q*_*B*_ = 0.0334369453801207. Fungicide parameters: *ω*_*A*_ = 0.5100900734551262, *θ*_*A*_ = 5.336781378462673, Λ_*A*_ = 0.003895561458581, *ω*_*B*_ = 0.8667335745106877, *θ*_*B*_ = 4.618154403228932, Λ_*B*_ = 0.0080007828566827.

**Supplementary Text 5 Figure 5.**
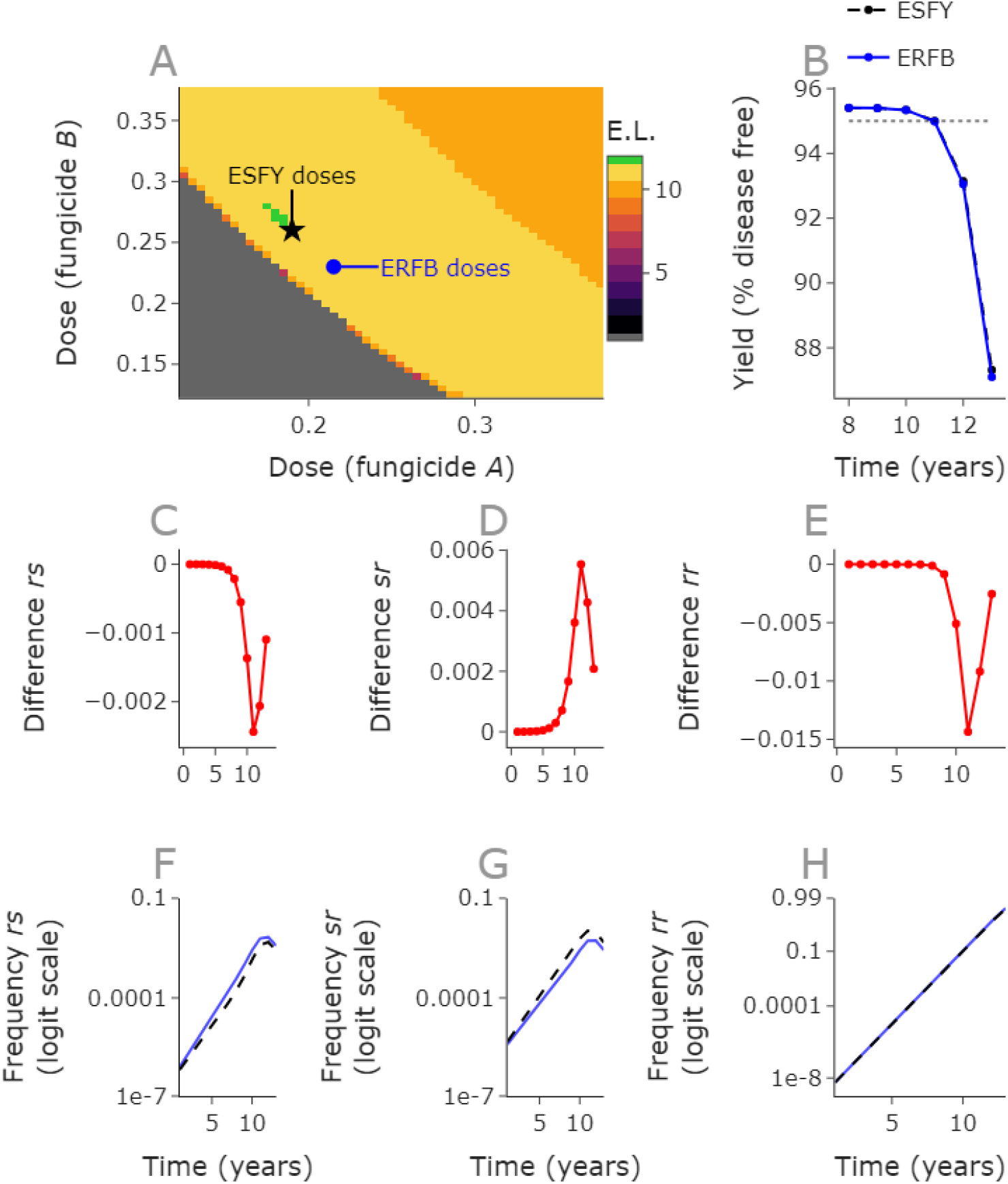
The single case where ESFY outperformed ERFB - explained. The optimal strategy achieves a yield of 95.002% in year 11, compared to the ERFB strategy which achieves 94.99% in this year (**B**). The optimal strategy fails in year 12, whereas the ERFB strategy fails in year 11 (**A**). In this case the ERFB strategy selects less strongly for the *rs* and *rr* strains but more strongly for the *sr* strain. **Parameter values:** Initial frequencies: *I*_*rr*_ = 1.3122930696727086 × 10^−10^, *I*_*rs*_ = 1.3687346286628245 × 10^−7^, *I*_*sr*_ = 9.216054320447214 × 10^−7^, *I*_*ss*_ = 0.9999989413898756. *q*_*B*_ = 0.0334369453801207. Fungicide parameters: *ω*_*A*_ = 0.5100900734551262, *θ*_*A*_ = 5.336781378462673, Λ_*A*_ = 0.003895561458581, *ω*_*B*_ = 0.8667335745106877, *θ*_*B*_ = 4.618154403228932, Λ_*B*_ = 0.0080007828566827, Doses: ESFY = 0.19, 0.26, ERFB = 0.215, 0.23.

